# Multi-plateau force-extension curves of long double-stranded DNA molecules

**DOI:** 10.1101/2023.03.12.532320

**Authors:** Alexander Y. Afanasyev, Alexey V. Onufriev

## Abstract

When highly stretched, double-stranded DNA exhibits a plateau region in its force-extension curve. Using a bead-spring coarse-grained dynamic model based on a non-convex potential, we predict that a long double-stranded DNA fragment made of several consecutive segments with substantially different plateau force values for each segment will exhibit multiple distinct plateau regions in the force-extension curve under physiologically relevant solvent conditions. For example, a long composite double-stranded (ds) DNA fragment consisting of two equal-length segments characterized by two different plateau force values, such as the poly(dA-dT)-poly(dG-dC) fragment, is predicted to exhibit two distinct plateau regions in its force-extension curve; a long composite dsDNA fragment consisting of three segments having three different plateau force values is predicted to have three distinct plateau regions. The formation of mixed states of slightly and highly stretched DNA, co-existing with macroscopically distinct phases of uniformly stretched DNA is also predicted.

When one of the segments overstretches, the extensions of the segments can differ drastically. For example, for the poly(dA-dT)-poly(dG-dC) composite fragment, in the middle of the first plateau, 96.7 % of the total extension of the fragment (relative to *L*_*x*_*/L*_0_ *≈* 1.0) comes from the poly(dA-dT) segment, while only 3.3 % of it comes from the poly(dG-dC) segment. The order of the segments has little effect on the force-extension curve or the distribution of conformational states. We speculate that the distinct structural states of stretched double-stranded DNA may have functional importance. For example, these states may modulate, in a sequence-dependent manner, the rate of double-stranded DNA processing by key cellular machines.

**TOC Graphic:** 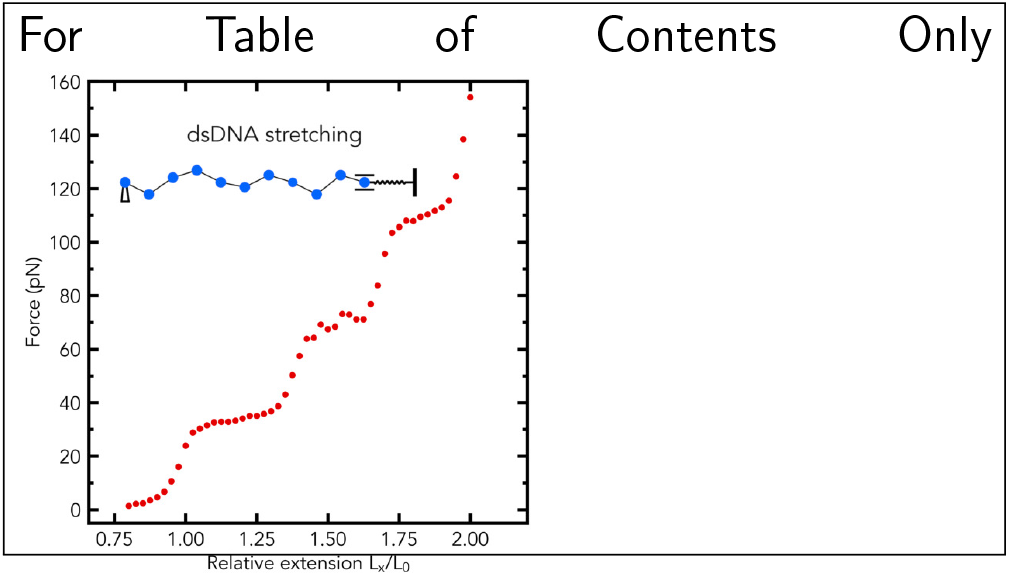

## Introduction

A double-stranded DNA (dsDNA) molecule is under bending, stretching, and torsion in living cells.^1–3^ Knowledge of the mechanical properties of the molecule^4^ is important to understanding of detailed mechanisms of many biological processes,^5–8^ as well as in designing of DNA-based nano-materials.^9^ Widely used experimental techniques employed to explore the mechanics of the polymer, the single-molecule stretching with optical tweezers and atomic force microscopy, reveal a peculiar distinct plateau region in force-extension curves of dsDNA.^6,10–18^

To describe force-extension curves of dsDNA from the single-molecule stretching experiments, including the distinct plateau region, a variety of theoretical models^19–26^ were proposed; a number of coarse-grained or fully atomistic molecular dynamics (MD) simulation studies revealed various aspects of the plateau region, including possible structural forms of the dsDNA.^27–37^ Sequence dependence of the mechanical properties of dsDNA was addressed in a number of computational^29,38–41^ and experimental works; ^14,15^ in particular, the plateau force values (the heights of the plateau) in the force-extension curves of dsDNA were found to be sequence dependent.^14,15,29^ Namely, under physiologically relevant solvent conditions, experiments show that the poly(dA-dT) DNA has a plateau force value of *∼* 35 pN,^15^ the poly(dG-dC) DNA has a plateau force value of *∼* 75 pN,^15^ and the torsionally constrained *λ*-phage DNA exhibits a plateau at *∼* 110 pN.^42^ However, to the best of our knowledge, experimental force-extension curves of long dsDNA fragments composed of any combination of long dsDNA segments with very distinct plateau values have not been reported.

So, here we ask a general question: what might happen if one were to stretch a composite dsDNA fragment made of sequentially connected multiple distinct DNA segments, under physiologically relevant conditions in an aqueous solution? Would such a long composite dsDNA fragment exhibit two or more distinct plateau regions in its force-extension curve? Likewise, how would the stretch of the entire fragment be distributed among the individual segments? For a set of sequentially connected harmonic springs of similar lengths and rigidity, the extension is distributed equally among them, but DNA is distinctly non-harmonic in the strong stretching regime, ^27^ suggesting the possibility of a very different scenario.

Specifically, we hypothesize that a long dsDNA fragment made up of several segments with distinctly different plateau force values will exhibit multiple plateau regions in the force-extension curve. We also hypothesize that a long dsDNA fragment composed of two long segments with two distinctly different plateau force values may show two distinct plateau regions in its force-extension curve. Similarly, we hypothesize that a long dsDNA fragment consisting of three long segments with three distinctly different plateau force values may show three distinct plateau regions in its force-extension curve.

Broadly speaking, there can be two non mutually exclusive, but distinct approaches to studying aspects of DNA deformation, including the specific questions posed above. One highly popular approach implies exploring, at the highest detail possible, the structures that emerge upon DNA deformation, including details of the conformational states^10,22,32,35,43–45^ that occur upon gradual deformation of the DNA double helix. On the other end of the spectrum are phenomenological models^19–21,23^ that aim at describing the general physics of DNA deformation, agnostic to the fine structural details. Some of these models can be highly coarse-grained; the main prerequisite for their success is that they rely on physically well-grounded effective potentials used to describe the interaction between the model particles. For example, the existence of the DNA overstretching plateau, or “softening” of strongly bent dsDNA, can be traced to the non-convexity of the corresponding potential functions, ^27,46^ the specific structural origins of the non-convexity do not matter for predictions that this model makes. Despite their obvious limitation – the absence of fine-grain description of the DNA structures seen along the deformation pathways – these models can still make valuable predictions about the overall behavior of the system, which tend to be robust to details. Here, we argue that it is this type of models that are best suited for the *initial* exploration of the questions posed above. Specifically, in this study, we computationally test our hypotheses by running coarse-grained MD simulations of long dsDNA fragments made up of several segments with distinctly different plateau force values. The distribution of the fragment extension between individual segments is analyzed in detail. We also explore whether the order of the distinct dsDNA segments that make up the whole fragments matters. Finally, the robustness of the predictions to the model details is confirmed.

### The Model. MD Simulations

We have performed Molecular Dynamics (MD) simulations of the stretching of dsDNA fragments using the previously developed bead-spring model, which is based on a non-convex stretching potential.^37^ The model reproduced the experimental stretching behavior of long dsDNA and dsRNA molecules, with one plateau region in their force-extension curves.^37^ Below are the key elements of the model,^37^ along with the full details of the MD simulations performed in this work.

All MD simulations of the stretching of DNA fragments were conducted in the ESPResSo 3.3.1 package^47^ with Langevin dynamics at *T* =300 Kelvin (K). The simulated system is composed of 100 beads. The bead size is set to 11 base pairs, which corresponds to one helical turn of B-DNA; the full rationale for this specific choice is given in Ref. ^37^ Briefly, the fact that the length of the minimal unit is much larger than a single base-pair is consistent with the findings of several experimental studies of the kinetics of the overstretching transition in *λ*-phage dsDNA,^43,48^ which reported the cooperative length of the transition from B-form to S-form (extended) is approximately 22-25 bp. In a sense, a cooperativity of the overstretching transition is “built into” the model, agnostic to details of the corresponding states at base-pair resolution. Each DNA molecule is modeled as a chain of *N* = 99 beads, connected by non-linear springs. The rightmost bead of the chain is connected to the right fixed end (i.e., 100th bead) via a linear spring. The linear spring represents the model of an optical trap. A schematic of the model, the model of the optical trap, and the applied boundary conditions are depicted in Figure 1A. Each bead of the chain represents an 11-bp segment of the DNA, with a diameter of 1 Å and a mass of 7150 Da. The stretching behavior of the non-linear spring is governed by the potential, *U* (*r*):

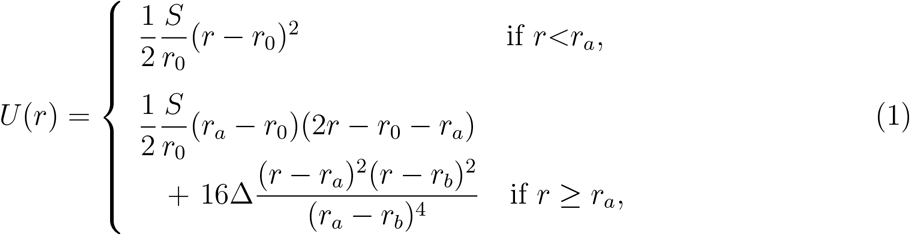

where *S* is the stretch modulus of the molecule (in *k*_*B*_*T* /Å), *r* is the distance between the centers of neighboring beads [the bond length (in Å)], *r*_0_ is the equilibrium bond length (in Å), *r*_*a*_ is the bond length at the beginning of the convex hull (in Å), *r*_*b*_ is the bond length at the end of the convex hull (in Å), and Δ is the energy difference between the potential *U* (in *k*_*B*_*T*) and the convex hull at *r*=(*r*_*a*_+*r*_*b*_)/2, as shown in Figure 1B. To account for the bending elasticity of the chain, characterized by its persistence length *P*, we use a discrete version of the worm-like chain (WLC) model.

**Figure 1:**
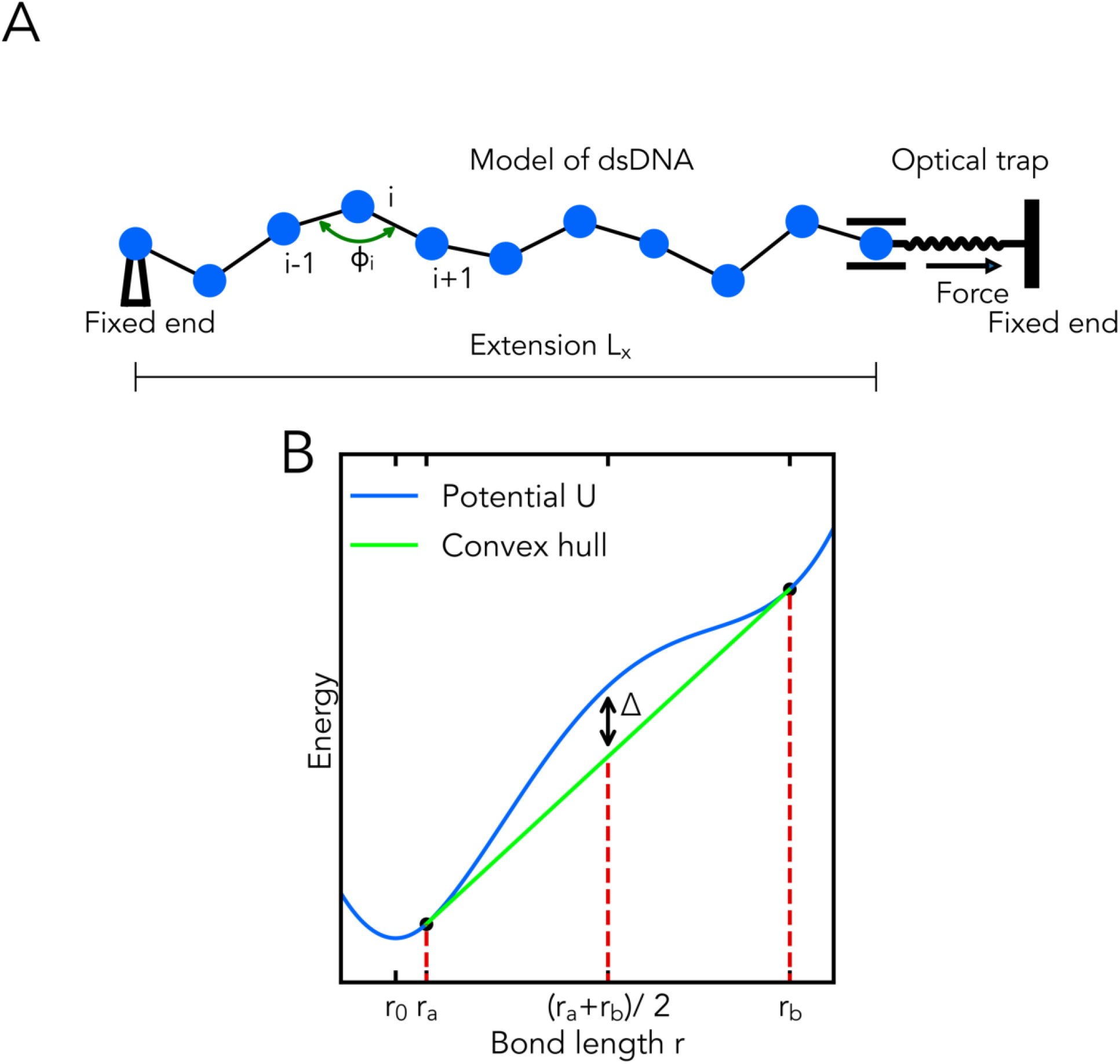
**(A)** Schematic showing the bead-spring model of the polymer chain in a random conformation, a model of the optical trap, and the applied displacement boundary conditions at the ends of the chain. The leftmost bead of the chain is fixed, and the rightmost bead is allowed to move in the horizontal direction only. **(B)** The non-convex potential energy *U* (*r*) (solid blue curve) governs the stretching behavior of the non-linear spring that connects each pair of beads. The existence of a nonconvex region in *U* (*r*), controlled by the parameter Δ, leads to nonuniform stretching of the chain.

The total potential energy of the chain, *E*_*chain*_, has two terms:

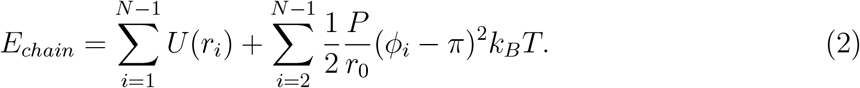

The first term represents the summation of the potential *U* over all adjacent pairs of beads, where *r*_*i*_ is the distance between the centers of bead *i* and bead (*i* + 1). The second term represents the harmonic bending energy of the chain (a discrete version of the worm-like chain (WLC) model), where *P* is its persistence length (in Å); *ϕ*_*i*_, the angle between the vectors from the centers of bead *i* to bead (*i−* 1) and bead (*i*+1), Figure 1A; the equilibrium bond length *r*_0_ is 11*h*, where the helical rise (i.e., the distance between two consecutive base pairs along the helical axis) *h* is set to 3.38 Å in all the simulations.

We model the optical trap as a harmonic spring potential *E*_*trap*_:

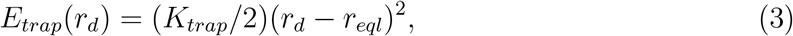

The corresponding force, *F*_*trap*_, is calculated as

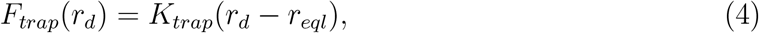

where the equilibrium length *r*_*eql*_ is 5*r*_0_, *r*_*d*_ is the distance between the right fixed end (fixed bead) and the center of the rightmost bead of the chain (Figure 1A), and *K*_*trap*_ is the stiffness of the optical trap. We make the following assumptions: (i) the average mass of a 1-bp segment of DNA is 650 Da, (ii) the contour length of dsDNA is fixed at *L*_0_ = (*N −* 1)*r*_0_, (iii) experimental force-extension curves were obtained at *T* =300 K, so *k*_*B*_*T ≈* 41.41 pN *·* Å.

The total potential energy of the system, *E*, is given as

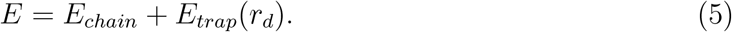

Unless stated otherwise, in all MD simulations the integration time step is 100 femtoseconds (fs), the duration of each simulation is 700 nanoseconds (ns), and each simulation starts from a conformation of the chain in which the beads are equally spaced, and in which the force (equation 4) is zero.

Unless stated otherwise, to obtain each point in the simulated force-extension curves, we pick the “one-phase” conformation, in which the beads are equally spaced with *r/r*_0_ = 0.8 as the starting conformation of the chain and in which the force (equation 4) is zero. The duration of each simulation is composed of a 50 ns equilibration run followed by a 650 ns production run. During the 650 ns production run, the values of the force in the optical trap and the relative extension *L*_*x*_*/L*_0_ are recorded at each integration time step and then averaged over 650 ns (6.5 million values). Note that since the stiffness of the linear spring (optical trap) is high, the amplitude of fluctuation of *L*_*x*_*/L*_0_ is relatively small during the simulation, i.e. for the starting conformation (the ‘one phase’) of the chain with *r/r*_0_ = 0.8, *L*_*x*_*/L*_0_ *≈* 0.8 at each time step. To obtain the next point in the force-extension curves, the above protocol is repeated starting from the same starting conformation, modified by a 0.025*r*_0_ increment to each bond length. The relative bond lengths are recorded every 0.1 ns during the 650 ns production run. This protocol aims to approximate an isometric experimental setup^a^, in which the end-to-end distance of the DNA molecule is kept fixed and the fluctuating force is measured.^49^ Since the stiffness of the optical trap *K*_*trap*_ (Figure 1A and equation 4) is much greater than the stiffness of the segments of the chain near the equilibrium bond length (table 2), the model approximates the isometric experiment reasonably closely, with the exact match corresponding to *K*_*trap*_ *→ ∞*.

In this work, the probability density *ρ* of relative bond lengths *r/r*_0_ is calculated with the *hist* function (density=True, bins=200) of Python’s Matplotlib Library. The “Bins” parameter defines the number of equal-width bins. If “density=True”, then the function normalizes the heights of bins so that the area under the histogram integrates to 1.

### dsDNA fragments used in the simulations

In this work, model polymer chains represent dsDNA fragments that are composed of combinations of specified dsDNA segments arranged sequentially. The following dsDNA segments are used: (i) torsionally unconstrained dsDNA of alternating A-T sequence (poly(dA-dT) segment) with a plateau force of *∼* 35 pN, (ii) torsionally unconstrained dsDNA of alternating G-C sequence (poly(dG-dC) segment) with a plateau force of *∼* 75 pN, and (iii) torsionally constrained *λ*-phage dsDNA (*λ*-DNA segment) with a plateau force of *∼* 110 pN. Each of the segments was previously explored experimentally and shown to exhibit a distinct plateau.^15,42^

We perform multiple MD simulations that model the stretching of the following dsDNA fragments: (1) the poly(dA-dT) fragment, (2) the poly(dG-dC) fragment, (3) the poly(dA-dT)-poly(dG-dC) fragment (Figure 2a), in which the first half of the fragment length is the poly(dA-dT) segment and the other half is the poly(dG-dC) segment, (4) the *λ*-DNA-poly(dG-dC) fragment^b^ (Figure 2b), in which the first half of the fragment length is the *λ*-DNA segment and the other half is the poly(dG-dC) segment, (5) the poly(dA-dT)-*λ*-DNA-poly(dG-dC) fragment (Figure 2c), in which the first *≈* 32.65 percent of the fragment length is the poly(dA-dT) segment, the second *≈* 33.67 percent of the fragment length is the *λ*-DNA segment, and the remaining *≈* 33.67 percent of the fragment length is the poly(dG-dC) segment and (6) dsDNA fragment composed of alternating poly(dA-dT) and poly(dG-dC) segments (alternating (poly(dA-dT)-poly(dG-dC)) fragment), Figure 2d.

**Figure 2:**
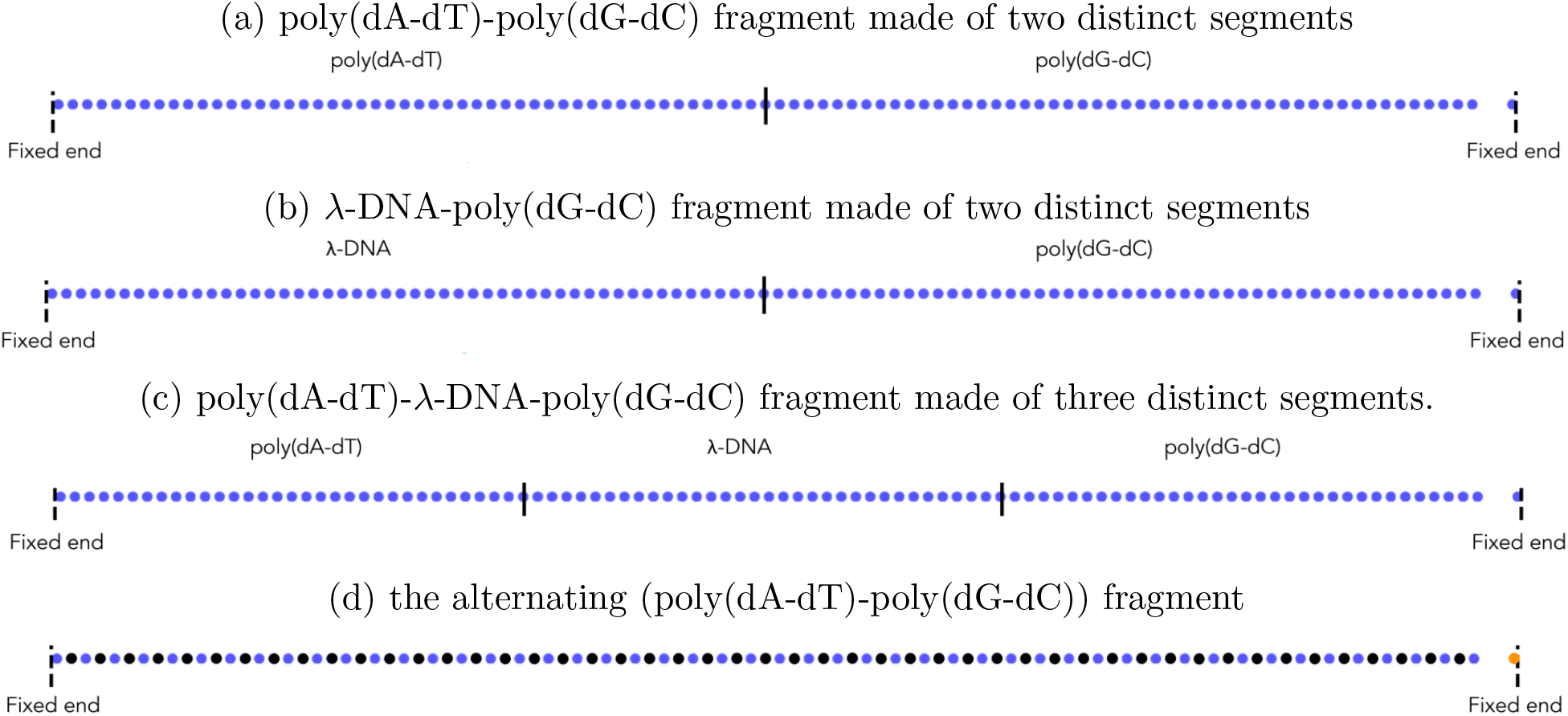
Model polymer chain representing the DNA fragment composed of several distinct segments as indicated. The solid vertical lines show the boundaries between the segments. Each dot is a coarse-grained bead, except for the rightmost dot that represents the right fixed end. In (d) the color of the dots (black – the poly(dA-dT) segment and blue – the poly(dG-dC) segment) indicates the segment between the dot of that color and its neighbor to the right. In this, and all other figures, the rightmost bead is for illustrative purposes only, its color does not carry any distance information.

The original experimental force-extension curves of the segments can be found in Figure 5 of Ref. 42, and Figures 9A and B of Ref. 15. In the case of the stretching of the alternating (poly(dA-dT)-poly(dG-dC)) fragment, we assume that its persistence length is 95.4 Å^37^.

The parameters of the model for the *λ*-DNA segment are shown in Figure 5A of Ref. 37, except for *r*_0_ which is 37.18 Å, assuming that the helical rise of the *λ*-DNA segment is 3.38 Å. We assume that running MD simulations with a small change in the value of *r*_0_ from 37.4 Å to 37.18 Å, while keeping all of the other parameters of the model the same, will lead to a simulated force-extension curve that is not very different from that of the *λ*-DNA segment, shown in Figure 5A of Ref. 37. In Tables 1 and 2, we summarize the parameters of the model for each segment and list other relevant simulation details.

**Table 1:** Parameters of the model for the poly(dA-dT) segment,^37^ the poly(dG-dC) segment,^37^ and the *λ*-DNA segment.

| Type | $r_0$<br>($\text{\AA}$) | $r_a/r_0$ | $r_b/r_0$ | $\Delta$<br>($k_B T$) | $S$<br>(pN) | $P$<br>($\text{\AA}$) |
| --- | --- | --- | --- | --- | --- | --- |
| poly(dA-dT) | 37.18 | 1.037 | 2.0 | 2 | 944 | 95.4 |
| poly(dG-dC) | 37.18 | 1.04 | 1.7 | 12 | 1864 | 41.9 |
| λ-DNA | 37.18 | 1.093 | 1.7 | 4 | 1191 | 616 |

**Table 2:** Simulation details for the poly(dA-dT) fragment, the poly(dG-dC) fragment, the poly(dA-dT)-poly(dG-dC) fragment, the *λ*-DNA-poly(dG-dC) fragment, the alternating (poly(dA-dT)-poly(dG-dC)) fragment, and the poly(dA-dT)-*λ*-DNA-poly(dG-dC) fragment; the Langevin collision frequency *γ* = 1*/T*_*p*_, where the period of oscillation of a bead with a mass *m* of 7150 Da attached to the spring (equation 1, if *r < r*_*a*_) 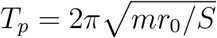; the stiffness of the optical trap *K*_*trap*_ = 10*S/r*_0_. For the composite fragments, *K*_*trap*_ and *γ* are determined by the parameters *S* and *r*_0_ of one of the segments. For integrator stability, we choose Δt *≪ T*_*p*_, where Δt is the integration time step.

| Type | $\gamma^a$<br>(ps<sup>-1</sup>) | $K_{trap}$<br>($k_B T / \text{\AA}^2$) |
| --- | --- | --- |
| poly(dA-dT) | 0.0232 | 6.129 |
| poly(dG-dC) | 0.0327 | 12.104 |
| poly(dA-dT)-poly(dG-dC) | 0.0232 | 6.129 |
| alternating (poly(dA-dT)-poly(dG-dC)) | 0.0232 | 6.129 |
| poly(dA-dT)-λ-DNA-poly(dG-dC) | 0.0261 | 7.733 |
| λ-DNA-poly(dG-dC) | 0.0232 | 6.129 |

<sup>a</sup> We deliberately use much smaller values of Langevin $\gamma$ compared to those of water to dramatically increase the speed of conformational sampling,<sup>50</sup> including the time needed to equilibrate the system. For long DNA fragments, the effective time scales that can be reached in this low $\gamma$ regime can be up to 100-fold the nominal trajectory length.<sup>50</sup> The small variation of $\gamma$ used for different segments has no tangible effect on the simulation outcomes on time scales much larger than $1/\gamma$ used to generate the production trajectories.

## Results and Discussion

### Two plateau regions

Our MD simulations show that two distinct plateau regions are indeed possible in the force-extension curve of the poly(dA-dT)-poly(dG-dC) fragment, Figure 3; this result can be compared to the familiar one-plateau force-extension curves for the poly(dA-dT) fragment or the poly(dG-dC) fragment, stretched individually (Figure 3). In the case of the poly(dA-dT)-poly(dG-dC) fragment, the segment with the lower plateau force, that is the poly(dA-dT) segment, overstretches first (i.e., within its plateau region), followed by the overstretching of the segment (the poly(dG-dC) segment) with the higher plateau force, as shown in Figure 3.

**Figure 3:**
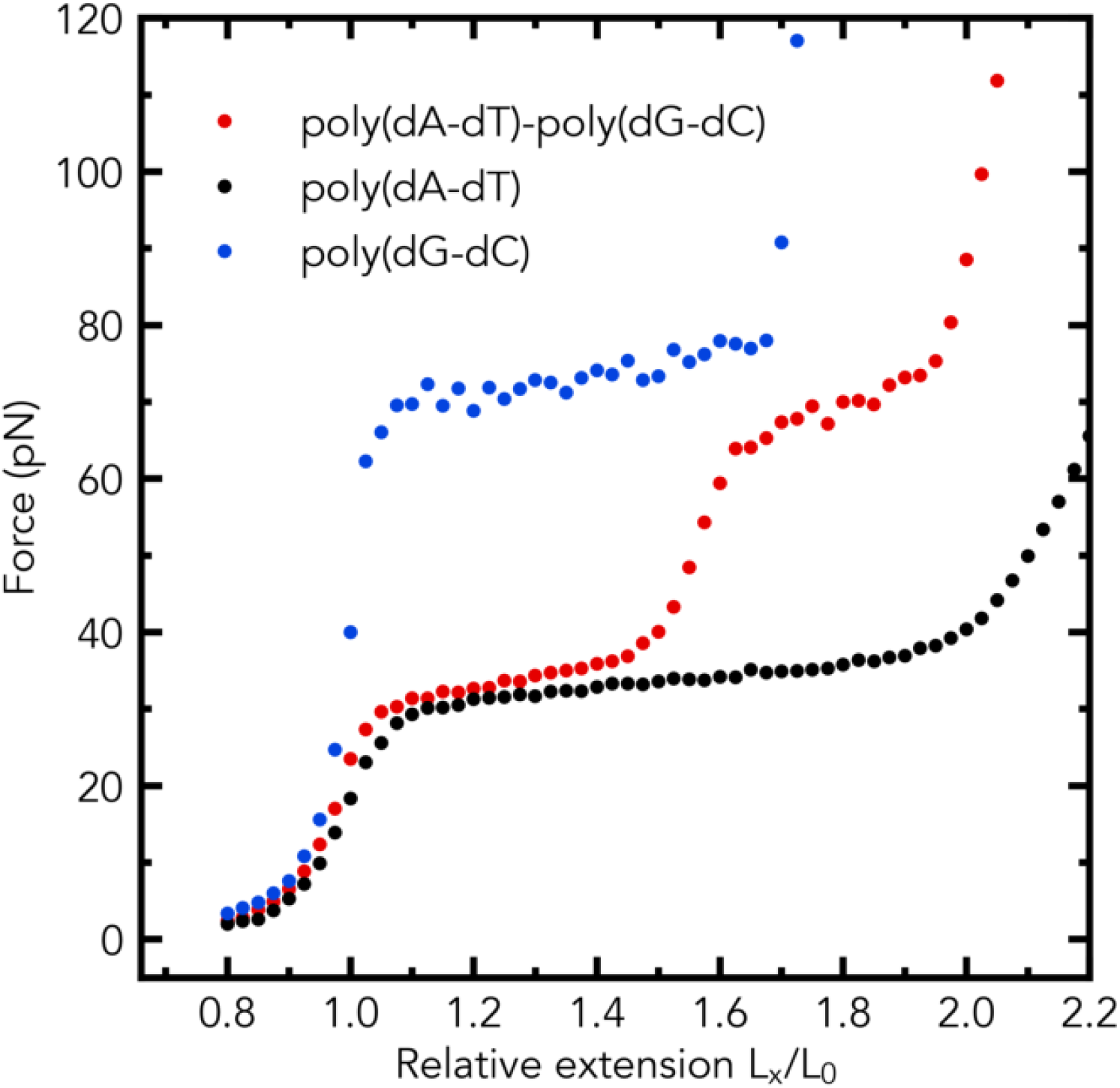
The emergence of two distinct plateau regions in the simulated force-extension curve of a dsDNA fragment made of two segments, Figure 2a, with distinctly different plateau force values. Two distinct plateau regions in the force-extension curve are predicted for the poly(dA-dT)-poly(dG-dC) fragment (1078 base pairs in total). Single distinct plateau regions in the force-extension curves (simulation) are shown for the poly(dA-dT) fragment and poly(dG-dC) fragment.

The distributions of the relative bond lengths *r/r*_0_ have two peaks in the middle of the first plateau region and three peaks in the middle of the second plateau region, see the insets of Figure 4. In the middle of the first plateau region of the force-extension curve, the simulation shows the formation of a macroscopically distinct phase in the poly(dG-dC) segment, but does not exhibit the formation of macroscopically distinct phases in the poly(dA-dT) segment, as illustrated in Figure 5. In the middle of the first plateau region, the poly(dA-dT) segment consists of a mix of two microscopic states: slightly and highly stretched. In contrast, in the middle of the second plateau region, the simulation shows the formation of a macroscopically distinct phase in the poly(dA-dT) segment, but does not show the formation of macroscopically distinct phases in the poly(dG-dC) segment. In the middle of the second plateau region, the poly(dG-dC) segment consists of a mix of two microscopic states: slightly and highly stretched, see the first two peaks in the top inset of Figure 4.

**Figure 4:**
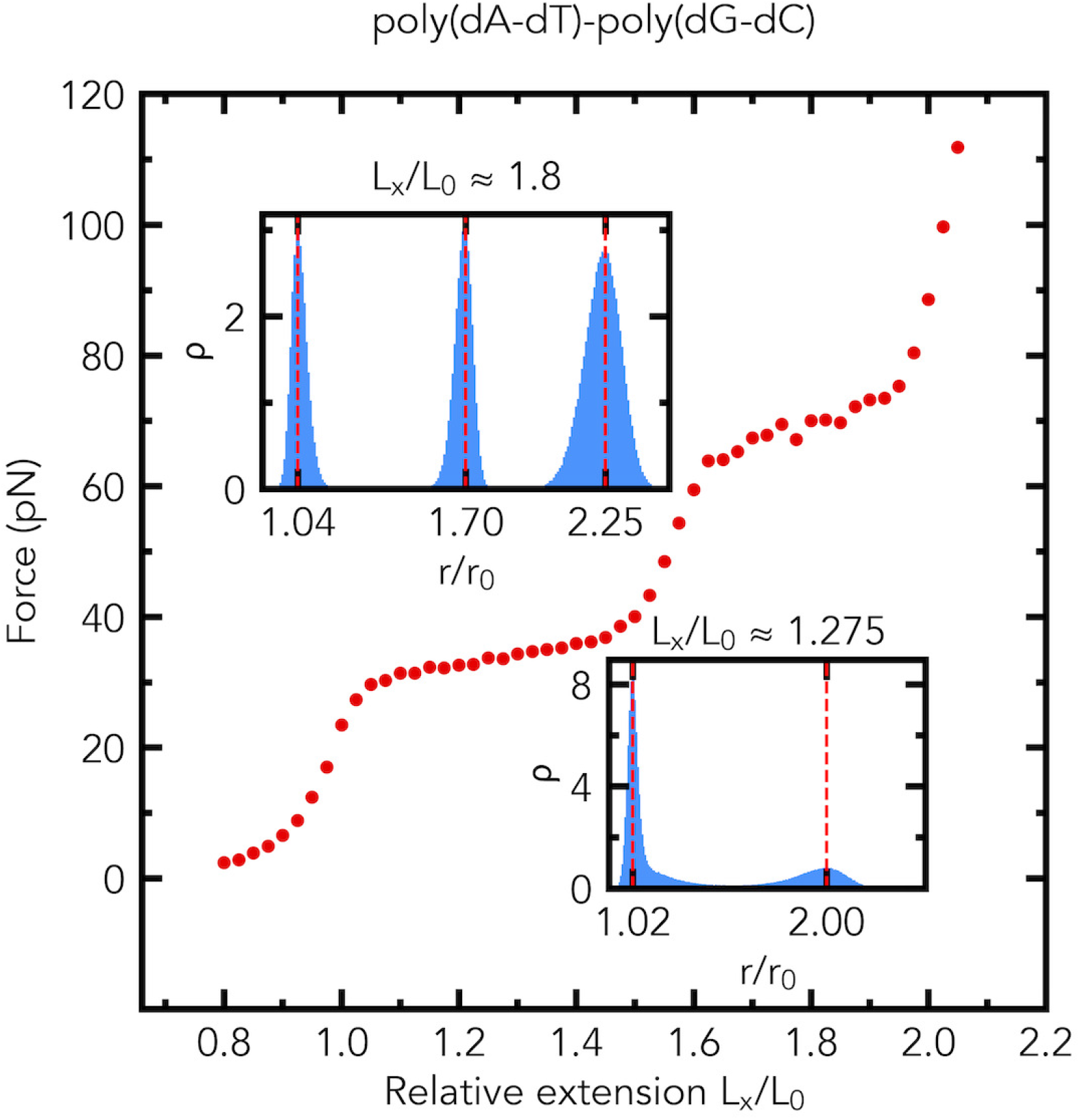
The distributions (the probability density *ρ*) of relative bond lengths *r/r*_0_ in the plateau regions of the simulated force-extension curve of a dsDNA fragment, Figure 2a, made of two segments with distinctly different plateau force values. Two plateau regions in the force-extension curve are predicted for the poly(dA-dT)-poly(dG-dC) fragment (1078 base pairs long). The average values of the force (in pN) at *L*_*x*_*/L*_0_ *≈* 1.275 and *L*_*x*_*/L*_0_ *≈* 1.8 are 33.6 *±* 0.4 (s.e.m., n=5) and 70 *±* 1.7 (s.e.m., n=5), respectively. Insets: the probability density (normalized histogram, number of bins is 200) *ρ* of relative bond lengths *r/r*_0_ at *L*_*x*_*/L*_0_ *≈* 1.275 (**Bottom**) and *L*_*x*_*/L*_0_ *≈* 1.8 (**Top**).

**Figure 5:**
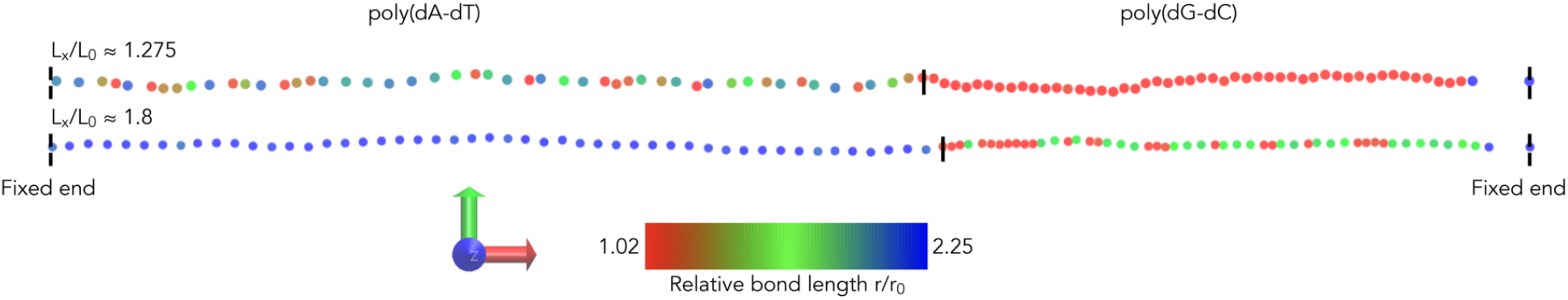
Coarse-grained conformations of the model poly(dA-dT)-poly(dG-dC) fragment in the plateau regions, at two values of the relative extension of the chain (**Top:** *L*_*x*_*/L*_0_ *≈* 1.275; **Bottom:** *L*_*x*_*/L*_0_ *≈* 1.8). The solid vertical lines show the boundaries between the poly(dA-dT) segment and the poly(dG-dC) segment. The color scale (from red – slight stretching, to blue – high stretching) indicates the relative bond length between the bead of that color and its neighbor to the right. The simulations start from the “one-phase” conformations, in which the beads are equally spaced with *r/r*_0_ = 1.275 (**Top**) and *r/r*_0_ = 1.8 (**Bottom**). Shown are the snapshots at t=700ns. Shown to the left of the color bar is a 3D Cartesian coordinate system. The red arrow is the x-direction, the green arrow is the y-direction, and the blue arrow indicates the z-direction.

Our MD simulations also show that two distinct plateau regions are present in the force-extension curve of the *λ*-DNA-poly(dG-dC) fragment, Figure 6. The distributions of the relative bond lengths *r/r*_0_ have two peaks in the middle of the first plateau region and in the middle of the second plateau region, see the insets of Figure 6. In the middle of the first plateau region of the force-extension curve, the simulation shows the formation of a macroscopically distinct phase in the *λ*-DNA segment, but does not exhibit the formation of macroscopically distinct phases in the poly(dG-dC) segment. In the middle of the first plateau region, the poly(dG-dC) segment consists of a mixture of two microscopic states: slightly and highly stretched. In contrast, in the middle of the second plateau region the simulation shows the formation of a macroscopically distinct phase in the poly(dG-dC) segment, but does not show the formation of macroscopically distinct phases in the *λ*-DNA segment. In the middle of the second plateau region, the *λ*-DNA segment consists of a mixture of two microscopic states: slightly and highly stretched.

**Figure 6:**
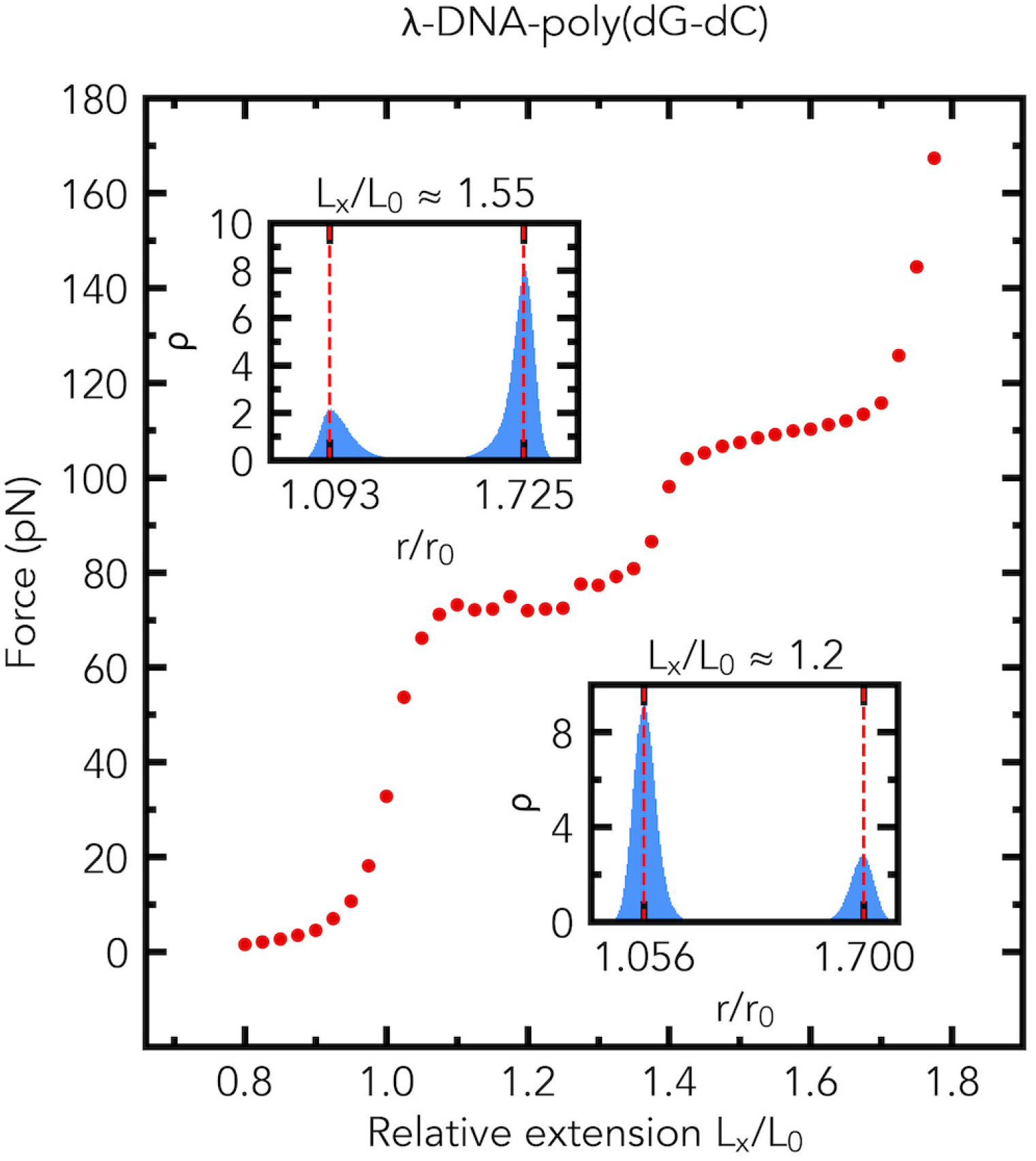
The emergence of two distinct plateau regions in the simulated force-extension curve of a dsDNA fragment made of two segments, Figure 2b, with distinctly different plateau force values. Two plateau regions in the force-extension curve are predicted for the *λ*-DNA-poly(dG-dC) fragment (1078 base pairs long). The distributions (the probability density *ρ*) of relative bond lengths *r/r*_0_ in the plateau regions are shown in the insets. The average values of force (in pN) at *L*_*x*_*/L*_0_ *≈* 1.2 and *L*_*x*_*/L*_0_ *≈* 1.55 are 72 *±* 1.5 (s.e.m., n=5) and 109.1 *±* 0.2 (s.e.m., n=5), respectively. Insets: the probability density (normalized histogram, number of bins is 200) *ρ* of relative bond lengths *r/r*_0_ at *L*_*x*_*/L*_0_ *≈* 1.2 (**Bottom**) and *L*_*x*_*/L*_0_ *≈* 1.55 (**Top**).

### Three plateau regions

Our MD simulations reveal that three distinct plateau regions can be possible in the force-extension curve of the poly(dA-dT)-*λ*-DNA-poly(dG-dC) fragment made of three distinct segments, Figure 7. The distributions of the relative bond lengths *r/r*_0_ have two peaks in the first plateau region and three peaks in the second and third plateau regions, see the insets of Figure 7. In the middle of the first plateau region, the simulation demonstrates the formation of a macroscopically distinct phase in the poly(dG-dC) segment and the *λ*-DNA segment, but does not show the formation of macroscopically distinct phases in the poly(dA-dT) segment, as illustrated in Figure 8. In the middle of the first plateau region, the poly(dA-dT) segment consists of a mix of two microscopic states: slightly and highly stretched. In contrast, in the middle of the second plateau region, the simulation reveals the formation of macroscopically distinct phases in the poly(dA-dT) segment and in the *λ*-DNA segment, but does not show the formation of macroscopically distinct phases in the poly(dG-dC) segment. In the middle of the second plateau region, the poly(dG-dC) segment consists of a mix of the two microscopic states: slightly and highly stretched. In the middle of the third plateau region, the simulation shows the formation of macroscopically distinct phases in the poly(dA-dT) segment and the poly(dG-dC) segment, but does not exhibit the formation of macroscopically distinct phases in the *λ*-DNA segment. In the middle of the third plateau region, the *λ*-DNA segment consists of a mix of two microscopic states: slightly and highly stretched, as shown in Figure 8. In this example the simulations start from the “one-phase” conformations, in which the beads are equally spaced with *r/r*_0_ = 1.175 (top), *r/r*_0_ = 1.525 (middle), and *r/r*_0_ = 1.85 (bottom). The corresponding time evolution of the chain in the plateau regions is illustrated by a series of snapshots shown in the SI.

**Figure 7:**
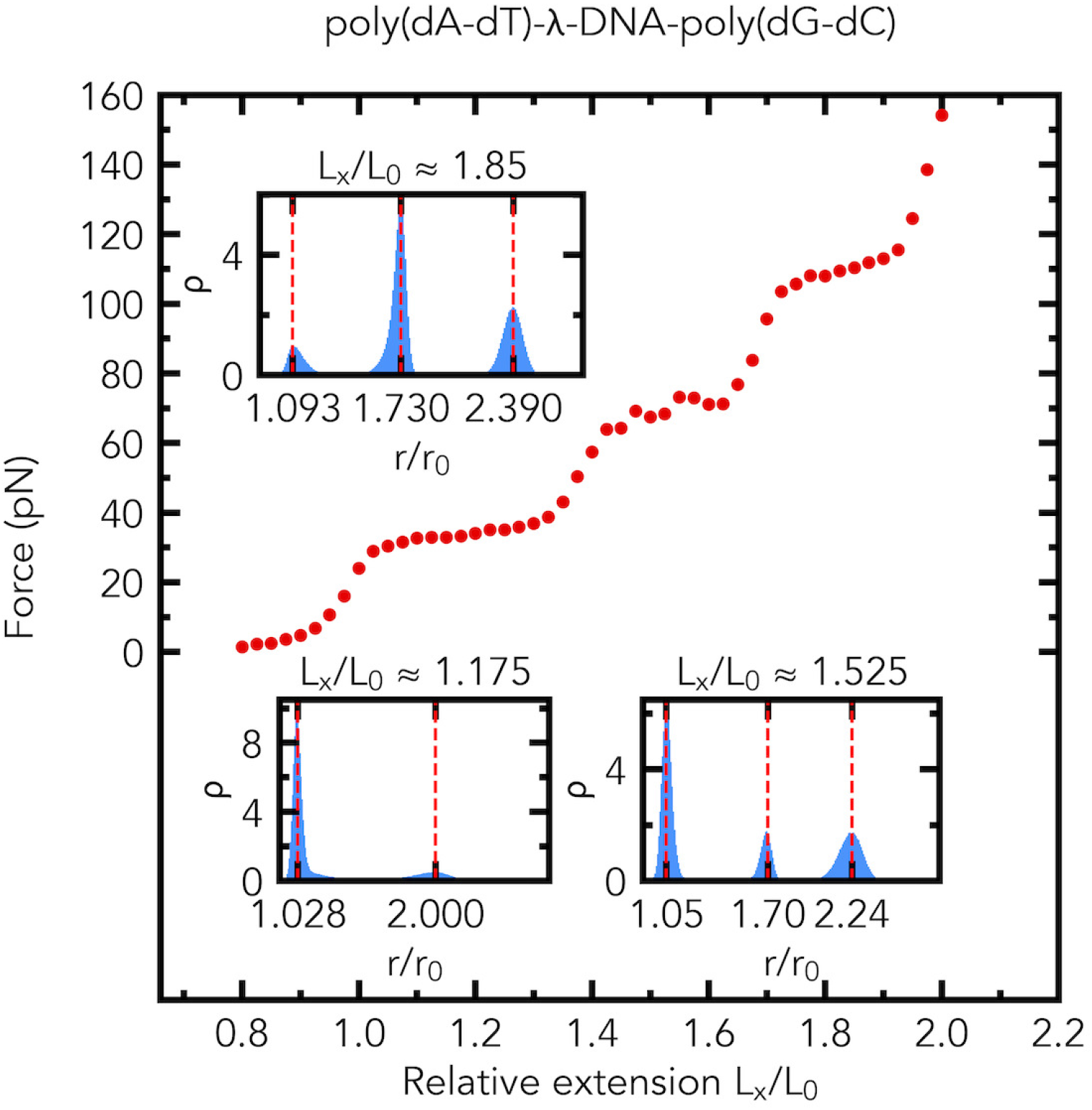
The emergence of three distinct plateau regions in the simulated force-extension curve of a dsDNA fragment made of three segments, Figure 2c, with distinctly different plateau force values. Three plateau regions in the force-extension curve are predicted for the poly(dA-dT)-*λ*-DNA-poly(dG-dC) fragment (1078 base pairs long). The corresponding distributions of relative bond lengths *r/r*_0_ in the plateau regions are shown in the insets. The average values of the force (in pN) at *L*_*x*_*/L*_0_ *≈*1.175, *L*_*x*_*/L*_0_ *≈*1.525, and *L*_*x*_*/L*_0_ *≈*1.85 are 33.2 *±*0.4 (s.e.m., n=5), 68.4*±*2.0 (s.e.m., n=5), and 110.3 *±*0.3 (s.e.m., n=5), respectively. Insets: the probability density (normalized histogram, number of bins is 200) *ρ* of relative bond lengths *r/r*_0_ at *L*_*x*_*/L*_0_ *≈* 1.175, *L*_*x*_*/L*_0_ *≈* 1.525, and *L*_*x*_*/L*_0_ *≈* 1.85.

**Figure 8:**
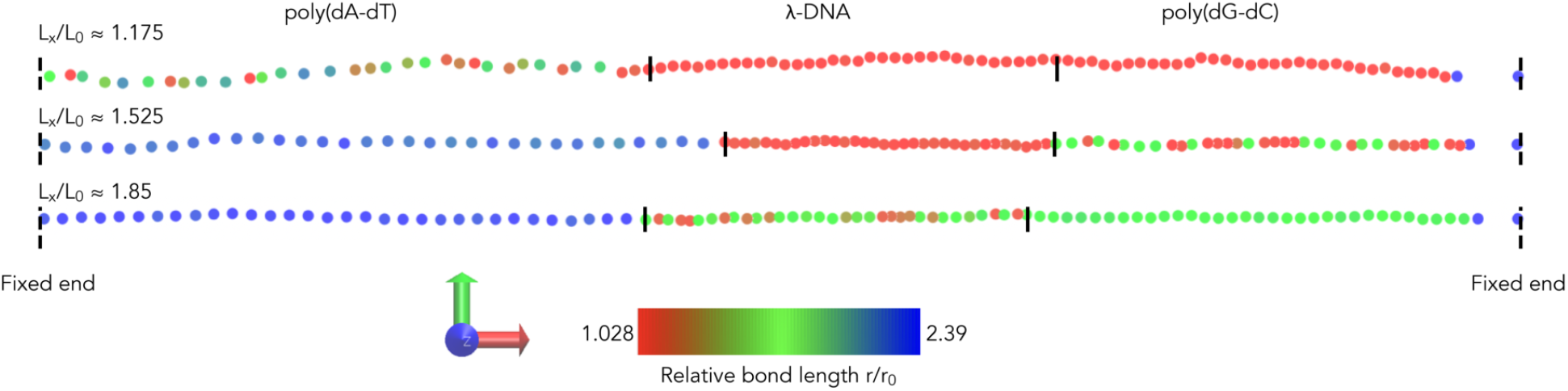
Coarse-grained conformations of the model poly(dA-dT)-*λ*-DNA-poly(dG-dC) fragment (at 700 ns) in the plateau regions, at three values of the relative extension of the chain (**Top:** *L*_*x*_*/L*_0_ *≈*1.175; **Middle:** *L*_*x*_*/L*_0_ *≈*1.525; **Bottom:** *L*_*x*_*/L*_0_ *≈*1.85). The solid vertical lines show the boundaries between the poly(dA-dT) segment, the *λ*-DNA segment, and the poly(dG-dC) segment. The color scale (from red – slight stretching, to blue – high stretching) indicates the relative bond length between the bead of that color and its neighbor to the right. The simulations start from the “one-phase” conformations, in which the beads are equally spaced with *r/r*_0_ = 1.175 (**Top**), *r/r*_0_ = 1.525 (**Middle**), and *r/r*_0_ = 1.85 (**Bottom**). Shown are the snapshots at t=700ns. Shown to the left of the color bar is a 3D Cartesian coordinate system. The red arrow is the x-direction, the green arrow is the y-direction, and the blue arrow indicates the z-direction.

### The extensions of the segments can be highly unequal

An interesting effect can be observed by examining the final equilibrated conformation of the chain of the poly(dA-dT)-poly(dG-dC) fragment in Figure 5. Namely, once the poly(dA-dT) segment is fully in the overstretching regime (middle of the first plateau region; *L*_*x*_*/L*_0_ *≈* 1.275), the poly(dG-dC) segment remains almost unstretched (i.e., *L*_*x*_*/L*_0_ *≈* 1.02). That is almost all the total stretch of the fragment, which is significant, is concentrated in the poly(dA-dT) segment. Quantitatively, at *L*_*x*_*/L*_0_ *≈* 1.0, the average end-to-end distance of the poly(dA-dT) segment is 1872 Å, and the average end-to-end distance of the poly(dG-dC) segment is 1771 Å. At *L*_*x*_*/L*_0_ *≈* 1.275, the average end-to-end distance of the poly(dA-dT) segment is 2840 Å, and the average end-to-end distance of the poly(dG-dC) segment is 1804 Å. Comparing the average end-to-end distances of the segments at *L*_*x*_*/L*_0_ *≈* 1.275 with those of *L*_*x*_*/L*_0_ *≈* 1.0, we conclude that 96.7 % of the extension of the fragment comes from the poly(dA-dT) segment, while only 3.3 % of it comes from the poly(dG-dC) segment.

There is a simple explanation for the phenomenon. Once the plateau region of any of the segments is reached, the stretching force pulling on the chain remains virtually constant, therefore those segments of the chain that are outside their respective overstretching regimes at this point gain virtually no additional extension, even though the entire chain is still being extended (at the “expense” of the segment currently in the plateau regime). This explanation is model-independent; therefore, we confidently predict that the effect will be observed whenever multiple distinct overstretching plateau regions are observed.

### The order of the segments has little effect

An interesting question arises whether the order in which various dsDNA segments are connected affects the resulting force-extension curves of the composite fragment, and/or the corresponding distributions of the relative bond lengths *r/r*_0_.

Our MD simulations show that two distinct plateau regions are observed in the force-extension curve of the alternating (poly(dA-dT)-poly(dG-dC)) fragment, Figure 9. As depicted in Figure 4 and Figure 9, the force-extension curve of the alternating (poly(dA-dT)-poly(dG-dC)) fragment is almost the same as that of the poly(dA-dT)-poly(dG-dC) fragment. Therefore, the segment order has little effect on the force-extension curves of the fragments. As in the case of the poly(dA-dT)-poly(dG-dC) fragment, the distributions of the relative bond lengths *r/r*_0_ have two peaks in the middle of the first plateau region and three peaks in the middle of the second plateau region, see the insets of Figure 9. Therefore, the segment order has little effect on the distributions of the relative bond lengths *r/r*_0_. In the first plateau region, the alternating (poly(dA-dT)-poly(dG-dC)) fragment consists of a mix of two microscopic states: slightly and highly stretched, whereas, in the middle of the second plateau region, the alternating (poly(dA-dT)-poly(dG-dC)) fragment is composed of a mix of three microscopic states. In the middle of the plateau regions of the force-extension curve, the simulations reveal no formation of distinct (macroscopic) phases in the model polymer chain for the alternating (poly(dA-dT)-poly(dG-dC)) fragment.

**Figure 9:**
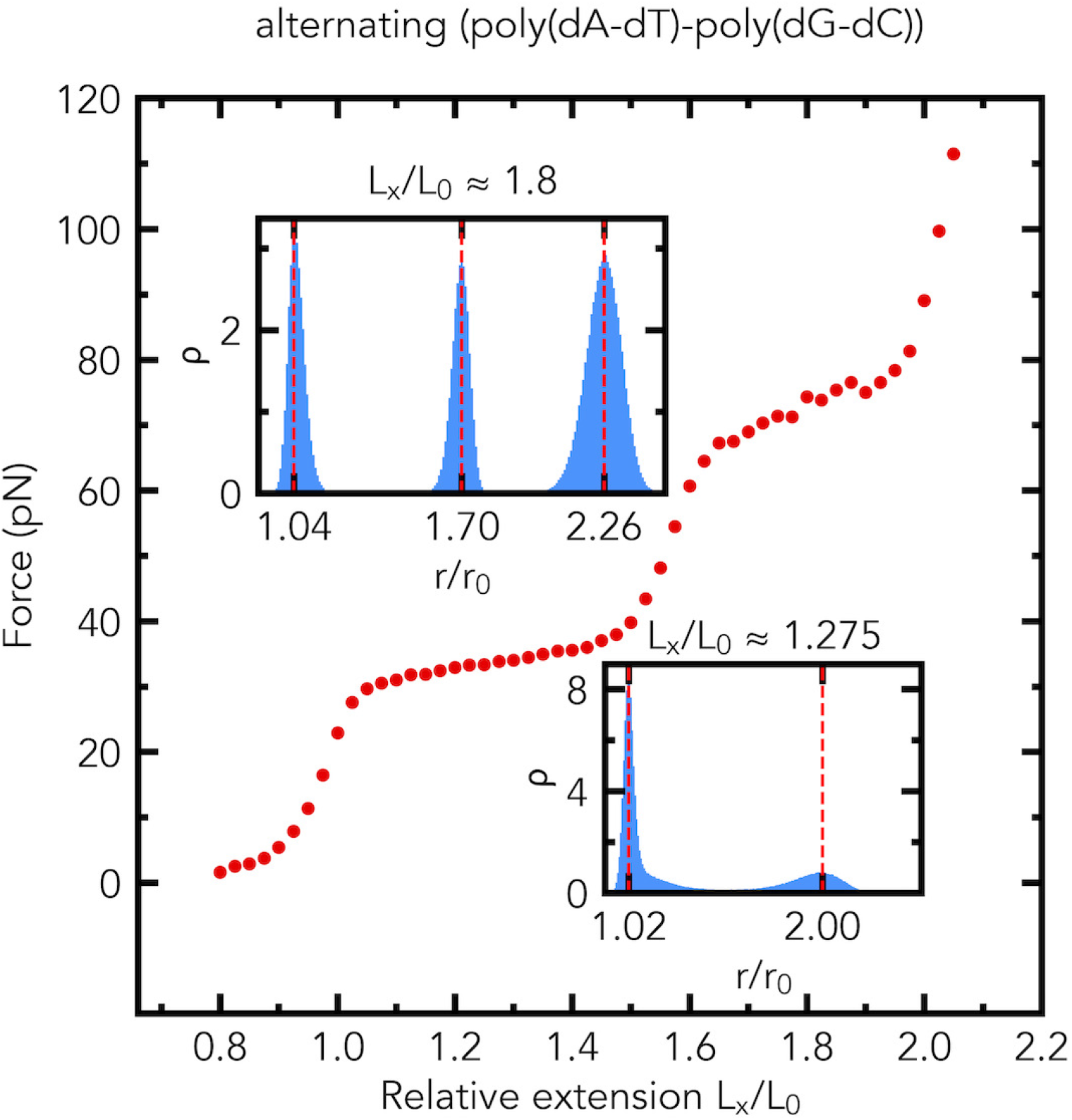
The segment order has little effect on the force-extension curve and distributions (the probability density *ρ*) of relative bond lengths *r/r*_0_. Shown are the simulated force-extension curve, and the distributions of relative bond lengths *r/r*_0_ of the alternating (poly(dA-dT)-poly(dG-dC)) fragment (1078 base pairs long), made of alternating poly(dA-dT) and poly(dG-dC) segments, Figure 2d. The force-extension curve and distributions (the probability density *ρ*) of relative bond lengths *r/r*_0_ are almost the same as those of the poly(dA-dT)-poly(dG-dC) fragment, Figure 4. The average values of the force (in pN) at *L*_*x*_*/L*_0_ *≈*1.275 and *L*_*x*_*/L*_0_ *≈*1.8 are 33.9*±* 0.1 (s.e.m., n=5) and 74.3 *±*0.9 (s.e.m., n=5), respectively. Insets: the probability density (normalized histogram, number of bins is 200) *ρ* of relative bond lengths *r/r*_0_ at *L*_*x*_*/L*_0_ *≈* 1.275 and *L*_*x*_*/L*_0_ *≈* 1.8.

### Peaks in the distribution of the relative bond lengths

In general, for a fragment that exhibits two distinct plateau regions, up to three peaks in the distributions of the relative bond lengths *r/r*_0_ may be expected, and for a fragment with three distinct plateau regions, as in Figure 7, up to four peaks may be expected. This is because, for a single segment, the distribution is bimodal at the plateau, and unimodal otherwise.^27,37^ However, whether two or three or even four peaks are observed depends on where the corresponding peaks occur in the individual segments, as two or three close peaks in different segments may give an appearance of one (broader) peak, when combined. We have verified the idea using the fragment in Figure 4. In the middle of the first plateau (*L*_*x*_*/L*_0_ *≈* 1.275) region of the force-extension curve of the poly(dA-dT)-poly(dG-dC) fragment, the distribution of the relative bond lengths *r/r*_0_ in the fragment shows only two distinct peaks. However, a detailed analysis of the distribution of the relative bond lengths *r/r*_0_ in the poly(dG-dC) and poly(dA-dT) segments individually (results not shown) demonstrates that the distribution in the poly(dG-dC) has one distinct peak (*r/r*_0_ = 1.02), the distribution in the poly(dA-dT) segment exhibits two distinct peaks (*r/r*_0_ = 1.037 and *r/r*_0_ = 2.0), but the one at *r/r*_0_ = 1.037 nearly coincides with the one at *r/r*_0_ = 1.02, resulting in the appearance of a bimodal distribution of the relative bond lengths *r/r*_0_ in the whole fragment.

## Conclusions

In this work, we have examined force-extension curves and the corresponding structural state of long dsDNA fragments consisting of shorter segments with distinct mechanical properties. Employing a bead-spring coarse-grained dynamic model based on a non-convex potential, we predict that a long double-stranded DNA fragment made up of several segments with different plateau force values for each segment will exhibit multiple distinct plateau regions in the force-extension curve under physiologically relevant solvent conditions. The order of the segments that make up the full DNA fragment has little effect on the force-extension curve or the distribution of the corresponding conformational states, at the resolution of the model (i.e., 11 base pairs long). However, to verify our predictions experimentally, we suggest that the distinct segments that make up the full dsDNA fragment should each be at least 100 base pairs long.

Our MD simulations reveal several specific examples where distinct multi-plateau regions are expected to appear in the force-extension curves of long composite dsDNA fragments under physiologically relevant solvent conditions. Specifically, our simulations show that a long dsDNA fragment comprising two equal-length segments with two different plateau force values, such as the poly(dA-dT)-poly(dG-dC) fragment, exhibits two distinct plateau regions. Three plateau regions are also possible in a fragment made up of three specific shorter segments. An example is a long DNA fragment consisting of three almost equal-length segments with three fairly different plateau force values In the case of the stretching of a long dsDNA fragment consisting of short, alternating poly(dA-dT) and poly(dG-dC) segments (the alternating (poly(dA-dT)-poly(dG-dC)) fragment, an equal fraction of each), our model reveals that two plateau regions are observed in the force-extension curve of the polymer without the formation of macroscopically distinct phases in the regions at room temperature. In the future, it will be interesting to explore how the lengths of the plateau regions of the force-extension curves depend on the fractions of the different sequences in the dsDNA fragments.

We also predict that when one of the segments making up the composite fragment over-stretches, the extensions of the segments can differ drastically. For example, in the middle of the first plateau region of the force-extension curve of the poly(dA-dT)-poly(dG-dC) fragment, 96.7 % of the total extension of the fragment (relative to *L*_*x*_*/L*_0_ *≈* 1.0) comes from the poly(dA-dT) segment, while only 3.3 % of it is from the poly(dG-dC) segment. A simple explanation for this phenomenon is proposed. The fact that the explanation is model-independent strongly suggests that the phenomenon is likely to be observed for other sequences for which multiple distinct overstretching plateau regions are observed.

The formation of mixed states of slightly and highly stretched DNA, co-existing with macroscopically distinct phases in several segments in the plateau regions, is also predicted. The formation of macroscopically distinct phases occurs in those segments that are not within their respective overstretching regions, so no separation into a two-state mixture is expected for those segments in the plateau regions. We speculate that the distinct structural states of stretched dsDNA may have functional importance. For example, these can modulate, in a sequence-dependent manner, the rate of dsDNA processing by key cellular machines.

We have proposed a straightforward criterion for constructing sequences that may lead to these unexplored regimes: the plateau force values of each segment making up the composite dsDNA fragment should differ substantially. This key condition is not particularly restrictive, which should facilitate choosing appropriate constructs for experimental verification, which may differ from those considered in this work. In particular, if constructing a dsDNA fragment fragment composed of a mix of the torsionally constrained and torsionally unconstrained segments proves difficult using standard biochemistry techniques, we would suggest considering an alternative: we predict that a long dsDNA fragment, composed of the poly(dA-dT) segment,^15^ the poly(dG-dC) segment,^15^ and the torsionally unconstrained *λ*-phage dsDNA segment of the same length (see, for example, Figure 3B of Ref. 37; the plateau force of *∼* 65 pN) might contain three plateau regions in its force-extension curve.

Similarly, two distinct plateau regions might be observed in the force-extension curve of a long dsDNA fragment consisting of the poly(dA-dT) ^15^ or poly(dG-dC) segment^15^ and the torsionally unconstrained *λ*-phage dsDNA segment (equal length each). The logic behind these predictions is simple: the segments that make up the composite fragment have quite different plateau force values. We speculate that single-molecule fluorescence resonance energy transfer, with judiciously placed labels, may be used to test the predictions regarding the nature of the structural states and their relative extensions that arise in multi-plateau regimes.

We are confident that our qualitative predictions are correct: the existence of two plateau regions was observed^10^ in the force-extension curve of *λ*-phage dsDNA in 5 M betaine solution. In the experiment, 5 M betaine may have enhanced the difference between the forces required to overstretch AT-rich and GC-rich regions of the experimental sequence, ^10^ leading to the observation of two plateau regions in the force-extension curve in this distinctly non-aqueous solvent. Specifically, 5 M betaine was thought to preferentially destabilize AT-rich regions over GC-rich regions.^10^ In our simulation, the enhancement is achieved differently, by choosing the sequences of the two DNA segments in a way to make their relevant differences large enough under physiological solution conditions, to yield two distinct plateau regions in the force-extension curve. Since our model is not designed to handle non-physiological solvent conditions, a direct comparison with the experimental data of Ref. 10 is not possible. It is also worth noting that our simulated double plateau force-extension curves of the poly(dA-dT)-poly(dG-dC) fragment and the torsionally constrained *λ*-phage dsDNA-poly(dG-dC) fragment qualitatively agree with a simulated force-extension curve predicted earlier by a multi-state statistical mechanical model for the strongly heterogeneous DNA sequence. ^51^

The main limitation of our approach is that it does not yet make predictions based on the DNA sequence alone: our model takes several relevant parameters as input, such as the value of the stretching force at the plateau. When accurate values of these parameters are readily available from the experiment, this limitation becomes a strength: we are highly confident in our predictions. On the other hand, some of the input parameters that came from the experiment had noticeable uncertainty. For example, the extension at which the plateau region ends in the experimental force-extension curve of the poly(dA-dT) fragment was difficult to extract from the experimental data, see Figure 7 of Ref. 37. In this work, we assumed that the extension at which the plateau region ends in the experimental force-extension curve of the poly(dA-dT) fragment is approximately two times the contour length of the poly(dA-dT) fragment (Figure 7 of Ref. 37). However, it is likely that the extension at which the plateau region ends in the experimental force-extension curve of the poly(dA-dT) fragment is approximately 1.7 times the contour length of the poly(dA-dT) fragment, which is on par with those of the torsionally constrained *λ*-phage dsDNA fragment, the torsionally unconstrained *λ*-phage dsDNA fragment, the poly(dG-dC) fragment. The above note is related to a related limitation of the model, which is that some of the experimental data points it relies on^14^ may have been updated. In the future, these updates may result in updates to the specific values of some of our quantitative predictions, such as the specific distribution of states, and the exact ratios of the relative segment extensions. We stress, however, that our key predictions here are qualitative, and these are robust to such details. These main qualitative predictions are, in fact, fairly simple, and follow from the general physical principles that our model is based on. It came to us as a surprise that these predictions had not been made explicit earlier.

## Supporting information

Supplemental Matherial

## Supporting Information Available

Supporting information contains a movie along with two figures illustrating the time evolution the poly(dA-dT)-*λ*-DNA-poly(dG-dC) fragment in the plateau regions at *L*_*x*_*/L*_0_ *≈* 1.175 and *L*_*x*_*/L*_0_ *≈* 1.85. It also contains an in-depth analysis of robustness of the results to various simulation details.

## Acknowledgement

We thank Jennifer S. Wayne and Romesh C. Batra for their helpful comments on the manuscript. This work was supported in part by the National Institutes of Health, grant R01 GM144596.

## AUTHOR DECLARATIONS

### Author Contributions

Alexander Y. Afanasyev: performed research and wrote the manuscript. Alexey V. Onufriev: designed research and wrote the manuscript.

## Footnotes

a Note that this setup is distinct from the alternative isotensional protocol, in which the force is held fixed and the fluctuating end-to-end distance is measured. However, it was shown ^49^ (based on a set of broad assumptions) that these two protocols result in similar force extension curves if the length of the DNA fragment is at least several times longer than its persistence length; in our simulations it is at least five times longer.

b While simulating a construct that combines torsionally constrained and unconstrained dsDNA within one fragment presents no problem to our coarse-grained model, studying force-extension properties of the same construct experimentally may be non-trivial. We use such constructs here only to illustrate what may happen when two segments with very distinct plateau force values are combined.

## References

(1) Ma, J.; Bai, L.; Wang, M. D. Transcription Under Torsion. Science 2013, 340, 1580– 1583.

(2) Burnham, D. R.; Kose, H. B.; Hoyle, R. B.; Yardimci, H. The mechanism of DNA unwinding by the eukaryotic replicative helicase. Nature Communications 2019, 10.

(3) Hall, M. A.; Shundrovsky, A.; Bai, L.; Fulbright, R. M.; Lis, J. T.; Wang, M. D. High-resolution dynamic mapping of histone-DNA interactions in a nucleosome. Nature Structural & Molecular Biology 2009, 16, 124–129.

(4) Marin-Gonzalez, A.; Vilhena, J. G.; Perez, R.; Moreno-Herrero, F. A molecular view of DNA flexibility. Quarterly Reviews of Biophysics 2021, 54.

(5) Lionnet, T.; Dawid, A.; Bigot, S.; Barre, F.-X.; Saleh, O. A.; Heslot, F.; Allemand, J.-F.; Bensimon, D.; Croquette, V. DNA mechanics as a tool to probe helicase and translocase activity. Nucleic Acids Research 2006, 34, 4232–4244.

(6) Basu, A.; Bobrovnikov, D. G.; Ha, T. DNA mechanics and its biological impact. Journal of Molecular Biology 2021, 433, 166861.

(7) Wasserman, M. R.; Liu, S. A Tour de Force on the Double Helix: Exploiting DNA Mechanics To Study DNA-Based Molecular Machines. Biochemistry 2019, 58, 4667– 4676.

(8) Lipfert, J.; Skinner, G. M.; Keegstra, J. M.; Hensgens, T.; Jager, T.; Dulin, D.; Kober, M.; Yu, Z.; Donkers, S. P.; Chou, F.-C. et al. Double-stranded RNA under force and torque: Similarities to and striking differences from double-stranded DNA. Proceedings of the National Academy of Sciences 2014, 111, 15408–15413.

(9) Li, N. K.; Kim, H. S.; Nash, J. A.; Lim, M.; Yingling, Y. G. Progress in molecular modelling of DNA materials. Molecular Simulation 2014, 40, 777–783, doi: 10.1080/08927022.2014.913792.

(10) Smith, S. B.; Cui, Y.; Bustamante, C. Overstretching B-DNA: The Elastic Response of Individual Double-Stranded and Single-Stranded DNA Molecules. Science 1996, 271, 795–799.

(11) van Mameren, J.; Gross, P.; Farge, G.; Hooijman, P.; Modesti, M.; Falkenberg, M.; Wuite, G. J. L.; Peterman, E. J. G. Unraveling the structure of DNA during over-stretching by using multicolor, single-molecule fluorescence imaging. Proceedings of the National Academy of Sciences 2009, 106, 18231–18236.

(12) Paik, D. H.; Perkins, T. T. Overstretching DNA at 65 pN Does Not Require Peeling from Free Ends or Nicks. Journal of the American Chemical Society 2011, 133, 3219–3221, PMID: 21207940.

(13) Herrero-Galán, E.; Fuentes-Perez, M. E.; Carrasco, C.; Valpuesta, J. M.; Carras-cosa, J. L.; Moreno-Herrero, F.; Arias-Gonzalez, J. R. Mechanical Identities of RNA and DNA Double Helices Unveiled at the Single-Molecule Level. Journal of the American Chemical Society 2013, 135, 122–131, PMID: 23214411.

(14) Rief, M.; Clausen-Schaumann, H.; Gaub, H. E. Sequence-dependent mechanics of single DNA molecules. Nat. Struct. Biol. 1999, 6, 346–349.

(15) Clausen-Schaumann, H.; Rief, M.; Tolksdorf, C.; Gaub, H. E. Mechanical Stability of Single DNA Molecules. Biophysical Journal 2000, 78, 1997–2007.

(16) King, G. A.; Gross, P.; Bockelmann, U.; Modesti, M.; Wuite, G. J. L.; Peterman, E. J. G. Revealing the competition between peeled ssDNA, melting bubbles, and S-DNA during DNA overstretching using fluorescence microscopy. Proceedings of the National Academy of Sciences 2013, 110, 3859–3864.

(17) King, G. A.; Peterman, E. J. G.; Wuite, G. J. L. Unravelling the structural plasticity of stretched DNA under torsional constraint. Nature Communications 2016, 7.

(18) Backer, A. S.; King, G. A.; Biebricher, A. S.; Shepherd, J. W.; Noy, A.; Leake, M. C.; Heller, I.; Wuite, G. J. L.; Peterman, E. J. G. Elucidating the Role of Topological Constraint on the Structure of Overstretched DNA Using Fluorescence Polarization Microscopy. The Journal of Physical Chemistry B 2021, 125, 8351–8361.

(19) Cizeau, P.; Viovy, J.-L. Modeling extreme extension of DNA. Biopolymers 1997, 42, 383–385.

(20) Ahsan, A.; Rudnick, J.; Bruinsma, R. Elasticity Theory of the B-DNA to S-DNA Transition. Biophysical Journal 1998, 74, 132–137.

(21) Storm, C.; Nelson, P. C. Theory of high-force DNA stretching and overstretching. Physical Review E 2003, 67.

(22) Léger, J. F.; Romano, G.; Sarkar, A.; Robert, J.; Bourdieu, L.; Chatenay, D.; Marko, J. F. Structural Transitions of a Twisted and Stretched DNA Molecule. Physical Review Letters 1999, 83, 1066–1069.

(23) Marko, J. F. DNA under high tension: Overstretching, undertwisting, and relaxation dynamics. Phys. Rev. E 1998, 57, 2134–2149.

(24) Haijun, Z.; Yang, Z.; Zhong-can, O.-Y. Bending and Base-Stacking Interactions in Double-Stranded DNA. Physical Review Letters 1999, 82, 4560–4563.

(25) Argudo, D.; Purohit, P. K. Equilibrium and Kinetics of DNA Overstretching Modeled with a Quartic Energy Landscape. Biophysical Journal 2014, 107, 2151–2163.

(26) Sarkar, A.; Léger, J.-F.; Chatenay, D.; Marko, J. F. Structural transitions in DNA driven by external force and torque. Physical Review E 2001, 63.

(27) Savin, A. V.; Kikot, I. P.; Mazo, M. A.; Onufriev, A. V. Two-phase stretching of molecular chains. Proceedings of the National Academy of Sciences 2013, 110, 2816–2821.

(28) Romano, F.; Chakraborty, D.; Doye, J. P. K.; Ouldridge, T. E.; Louis, A. A. Coarse-grained simulations of DNA overstretching. The Journal of Chemical Physics 2013, 138, 085101.

(29) Luan, B.; Aksimentiev, A. Strain Softening in Stretched DNA. Phys. Rev. Lett. 2008, 101, 118101.

(30) Fiasconaro, A.; Falo, F. Dynamical model for the full stretching curve of DNA. Phys. Rev. E 2012, 86, 032902.

(31) Pupo, A. E. B.; Falo, F.; Fiasconaro, A. DNA overstretching transition induced by melting in a dynamical mesoscopic model. The Journal of Chemical Physics 2013, 139, 095101.

(32) Cluzel, P.; Lebrun, A.; Heller, C.; Lavery, R.; Viovy, J.-L.; Chatenay, D.; Caron, F. DNA: An Extensible Molecule. Science 1996, 271, 792–794.

(33) Lebrun, A.; Lavery, R. Modelling Extreme Stretching of DNA. Nucleic Acids Research 1996, 24, 2260–2267.

(34) Li, H.; Gisler, T. Overstretching of a 30 bp DNA duplex studied with steered molecular dynamics simulation: Effects of structural defects on structure and force-extension relation. The European Physical Journal E 2009, 30.

(35) Luan, B.; Aksimentiev, A. Strain Softening in Stretched DNA. Physical Review Letters 2008, 101.

(36) Wolter, M.; Elstner, M.; Kubař, T. On the Structure and Stretching of Microhydrated DNA. The Journal of Physical Chemistry A 2011, 115, 11238–11247.

(37) Afanasyev, A. Y.; Onufriev, A. V. Stretching of Long Double-Stranded DNA and RNA Described by the Same Approach. Journal of Chemical Theory and Computation 2022, 18, 3911–3920.

(38) Marin-Gonzalez, A.; Vilhena, J. G.; Moreno-Herrero, F.; Perez, R. Sequence-dependent mechanical properties of double-stranded RNA. Nanoscale 2019, 11, 21471–21478.

(39) Marin-Gonzalez, A.; Vilhena, J.; Moreno-Herrero, F.; Perez, R. DNA Crookedness Regulates DNA Mechanical Properties at Short Length Scales. Physical Review Letters 2019, 122.

(40) Lankaš, F.; Šponer, J.; Hobza, P.; Langowski, J. Sequence-dependent elastic properties of DNA. Journal of Molecular Biology 2000, 299, 695–709.

(41) Lankaš, F. DNA sequence-dependent deformability-insights from computer simulations. Biopolymers 2004, 73, 327–339.

(42) King, G. A.; Burla, F.; Peterman, E. J. G.; Wuite, G. J. L. Supercoiling DNA optically. Proceedings of the National Academy of Sciences 2019, 116, 26534–26539.

(43) Bianco, P.; Bongini, L.; Melli, L.; Dolfi, M.; Lombardi, V. PicoNewton-Millisecond Force Steps Reveal the Transition Kinetics and Mechanism of the Double-Stranded DNA Elongation. Biophysical Journal 2011, 101, 866–874.

(44) Bongini, L.; Lombardi, V.; Bianco, P. The transition mechanism of DNA overstretching: a microscopic view using molecular dynamics. Journal of The Royal Society Interface 2014, 11, 20140399.

(45) Schakenraad, K.; Biebricher, A. S.; Sebregts, M.; ten Bensel, B.; Peterman, E. J. G.; Wuite, G. J. L.; Heller, I.; Storm, C.; van der Schoot, P. Hyperstretching DNA. Nature Communications 2017, 8, 2197.

(46) Drozdetski, A. V.; Mukhopadhyay, A.; Onufriev, A. V. Strongly Bent Double-Stranded DNA: Reconciling Theory and Experiment. Frontiers in Physics 2019, 7, 195.

(47) Limbach, H.; Arnold, A.; Mann, B.; Holm, C. ESPResSo—an extensible simulation package for research on soft matter systems. Computer Physics Communications 2006, 174, 704–727.

(48) Bongini, L.; Melli, L.; Lombardi, V.; Bianco, P. Transient kinetics measured with force steps discriminate between double-stranded DNA elongation and melting and define the reaction energetics. Nucleic Acids Research 2013, 42, 3436–3449.

(49) Keller, D.; Swigon, D.; Bustamante, C. Relating Single-Molecule Measurements to Thermodynamics. Biophysical Journal 2003, 84, 733–738.

(50) Anandakrishnan, R.; Drozdetski, A.; Walker, R.; Onufriev, A. Speed of Conformational Change: Comparing Explicit and Implicit Solvent Molecular Dynamics Simulations. Biophysical Journal 2015, 108, 1153–1164.

(51) Whitelam, S.; Pronk, S.; Geissler, P. L. Stretching chimeric DNA: A test for the putative S-form. The Journal of Chemical Physics 2008, 129, 205101.

