## Supplemental Matherial for "Multi-plateau force-extension curves of long double-stranded DNA molecules"

### Supplementary Material For: Multi-plateau force-extension curves of long double-stranded DNA molecules

§ *Present address: Department of Chemistry, The University of Texas at Austin, 105 E 24TH ST, Austin, TX 78712, USA*

#### Time evolution of poly(dA-dT)-$\lambda$-DNA-poly(dG-dC) fragment

Time evolution of the chain for the poly(dA-dT)-$\lambda$-DNA-poly(dG-dC) fragment in the plateau regions at $L_x/L_0 \approx 1.175$ and $L_x/L_0 \approx 1.85$ is illustrated in Figures S1 and S2, respectively.

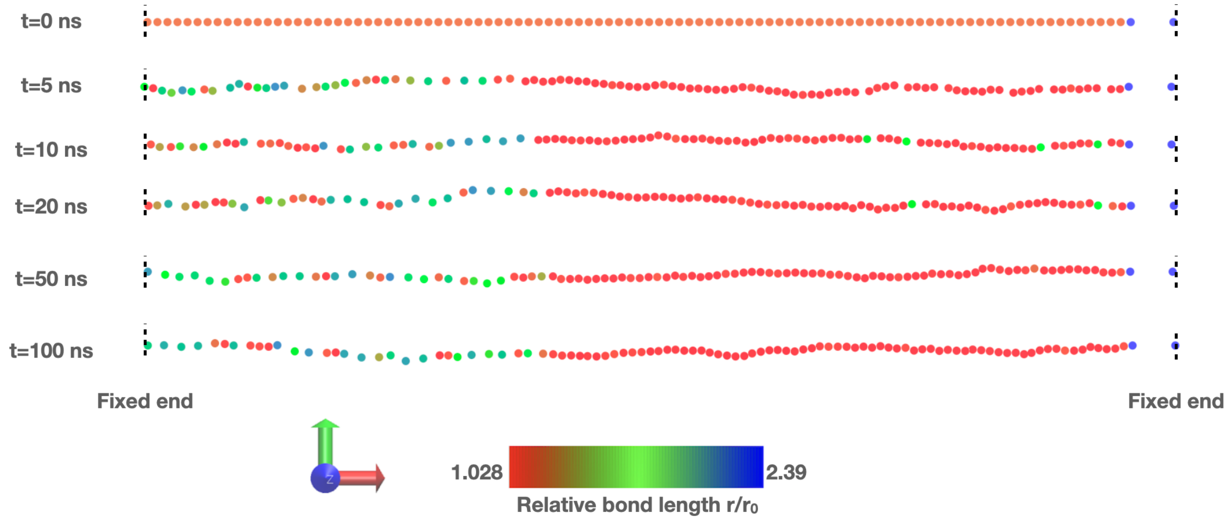


Figure S1: Time evolution of the chain for the poly(dA-dT)-$\lambda$-DNA-poly(dG-dC) fragment in the plateau region at $L_x/L_0 \approx 1.175$. The color scale (from red – slight stretching, to blue – high stretching) indicates the relative bond length between the bead of that color and its neighbor to the right. The simulation starts from the "one-phase" conformation, in which the beads are equally spaced with $r/r_0 = 1.175$.

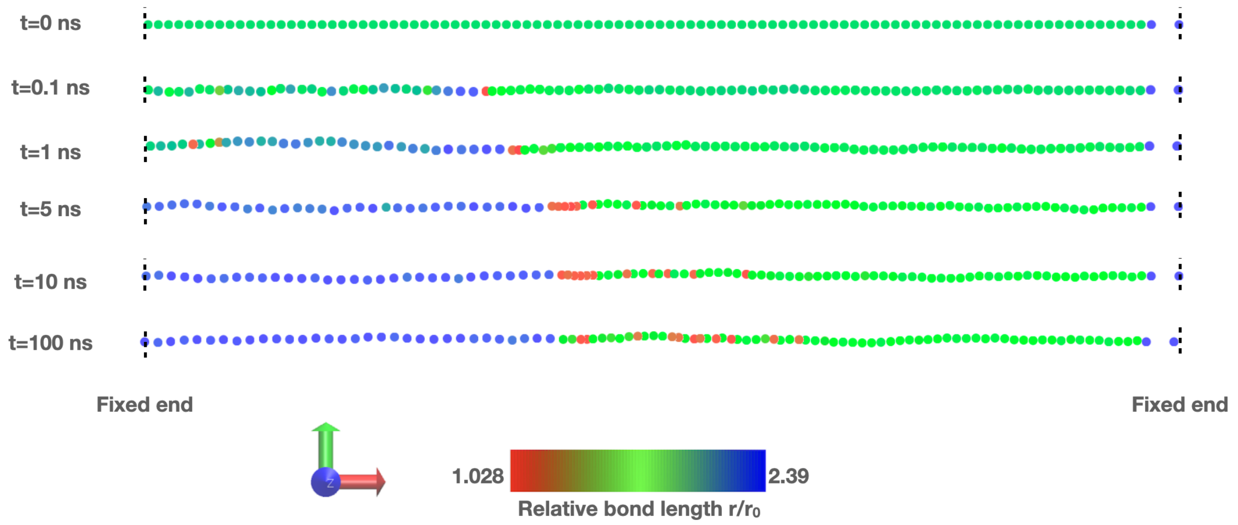


Figure S2: Time evolution of the chain for the poly(dA-dT)-$\lambda$-DNA-poly(dG-dC) fragment in the plateau region at $L_x/L_0 \approx 1.85$. The color scale (from red – slight stretching, to blue – high stretching) indicates the relative bond length between the bead of that color and its neighbor to the right. The simulation starts from the "one-phase" conformation, in which the beads are equally spaced with $r/r_0 = 1.85$. The corresponding movie is available as a supplemental material.

#### Robustness of results with respect to simulation details

##### Simulation length

To check whether a simulation time of 700 ns is long enough, we increased the simulation time from 700 ns to 2000 ns for the poly(dA-dT)-$\lambda$-DNA-poly(dG-dC) fragment. Figure S3 shows that a near tripling of the simulation time has little effect on the force-extension curve, and the distributions (the probability density $\rho$) of relative bond lengths $r/r_0$ in the plateau regions. As shown in Figure S3, the average values of the force (equation 4 of the main text) obtained from the shorter simulation may deviate by about $\sim 10$ % from the longer run result in the plateau region, where the poly(dG-dC) segment consists of a mix of two states: slightly and highly stretched. We consider this deviation acceptable for the main conclusions made in this work. In Figure S3, the simulations start from the “one-phase” conformations in which the beads are equally spaced.

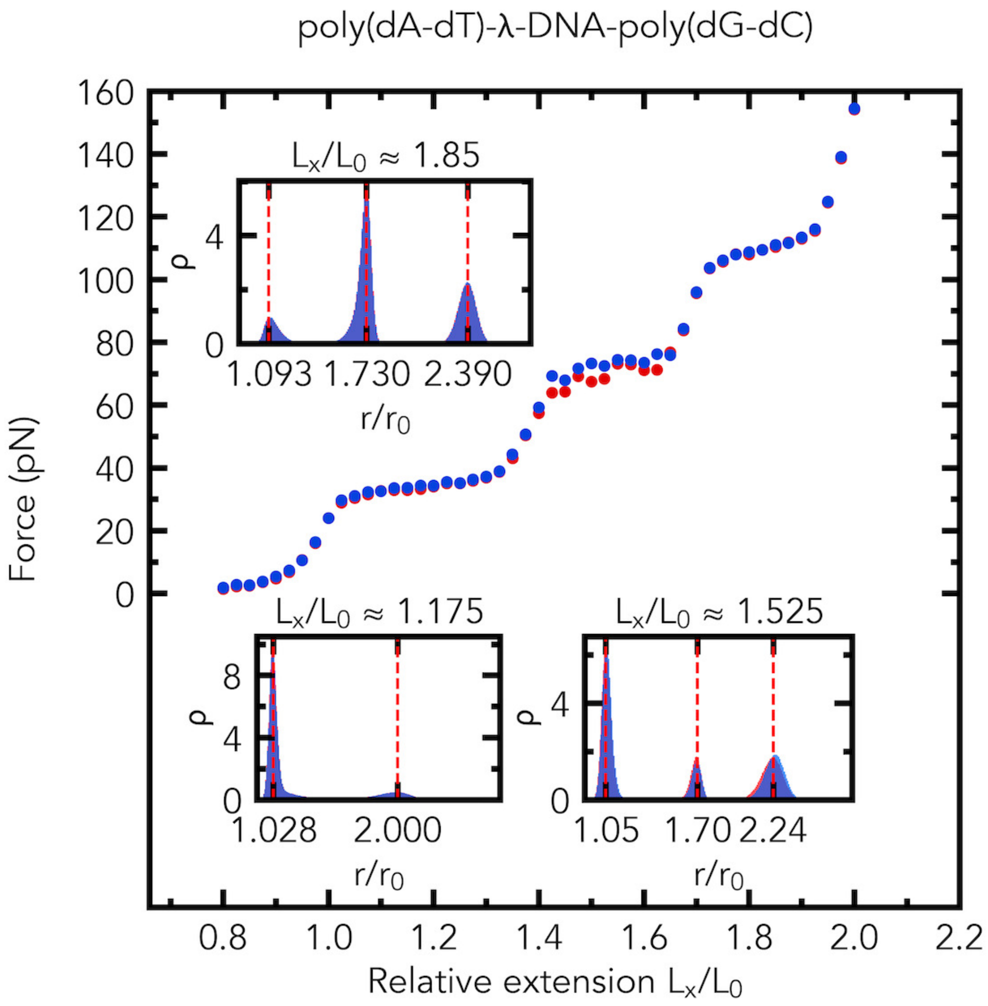


Figure S3: Robustness of results with respect to simulation time for the poly(dA-dT)-$\lambda$-DNA-poly(dG-dC) fragment. The values of the force (equation 4 of the main text) and relative extension are recorded at each integration time step, and then averaged over either the last 650 ns (6.5 million values) of the 700 ns simulation run (red dots), or over the last 650 ns (6.5 million values) of the 2000 ns simulation run (blue dots). Insets: the probability density (normalized histogram, number of bins is 200) $\rho$ of relative bond lengths $r/r_0$ at $L_x/L_0 \approx 1.175$, $L_x/L_0 \approx 1.525$, and $L_x/L_0 \approx 1.85$. In the insets: red data are for a simulation time of 700 ns, and blue data are for a simulation time of 2000 ns; the relative bond lengths are recorded every 0.1 ns of the last 650 ns of the simulation run.

Also, Figure S4 shows how the potential energy of the system changes with time at $L_x/L_0 \approx 1.175$. After about 25 ns of simulation, the potential energy begin to fluctuate about its average value. We can consider this behavior as a sign that, with respect to the total potential energy, the system approaches an equilibrium for the poly(dA-dT)-$\lambda$-DNA-poly(dG-dC) fragment at $L_x/L_0 \approx 1.175$. In Figure S4, the simulation starts from the "one-phase" conformation, in which the beads are equally spaced with $r/r_0 = 1.175$.

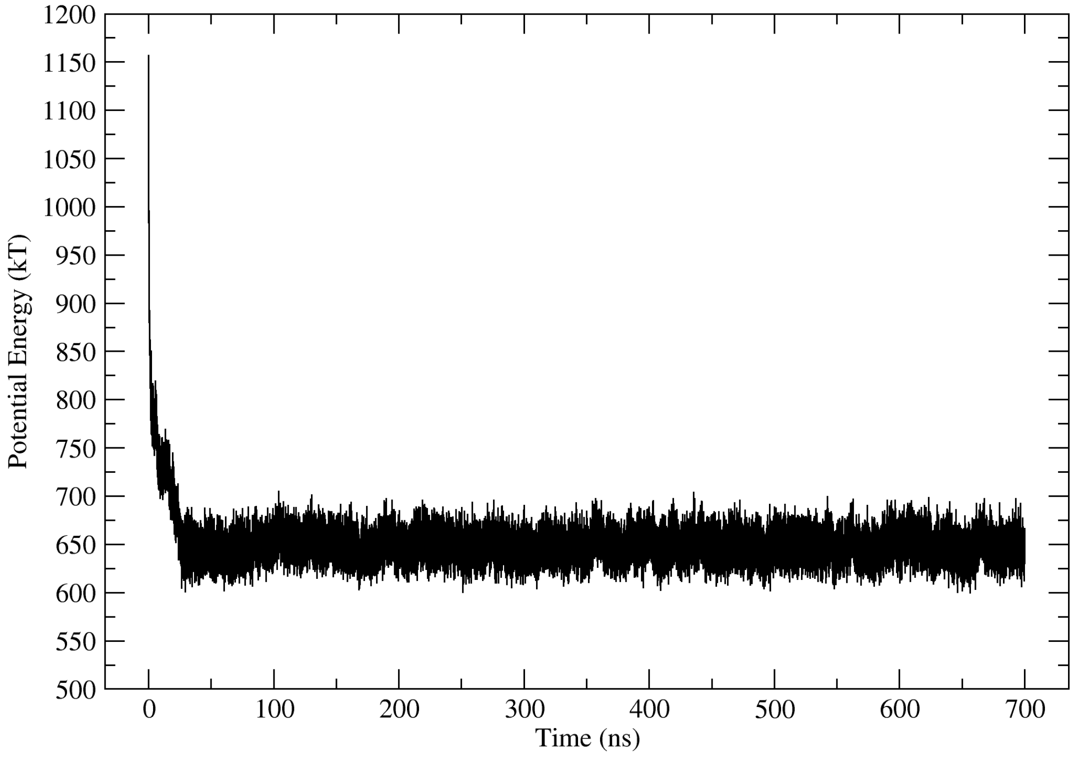


Figure S4: Potential energy of the system vs. time at $L_x/L_0 \approx 1.175$ for the poly(dA-dT)-$\lambda$-DNA-poly(dG-dC) fragment. The system contains the model of the optical trap that is represented as a linear spring. The stiffness of the linear spring is much higher than those of the non-linear springs that connect the chain, so we can conclude that the contribution of the potential energy of the linear spring to the total potential energy of the system is small. Therefore, the total potential energy of the system is approximately equal to the total potential energy of the chain.

##### Long-range interactions

To verify the robustness of our predictions with respect to the presence of long-range electrostatic interactions in the model, we added weak long-range electrostatic interactions explicitly (the Debye-Hückel potential) to the model for the poly(dA-dT)-$\lambda$-DNA-poly(dG-dC) fragment. The Debye-Hückel potential, $U_{DH}(r) = l_b k_B T q^2 \exp(-\kappa r)/re^2$, is added to explicitly model long-range interactions between charged beads in a salt solution for $r < r_c$, where $r$ is the distance between the centers of two beads, $e$ is the elementary charge, the Bjerrum length $l_b$ is 10 Å, the inverse Debye length $\kappa$ is 0.01 Å$^{-1}$, and the force cutoff $r_c$ is $10r_0$. Since the model already includes the long-range electrostatic interactions implicitly, via its parameters such as the stretch modulus and the persistence length, we set the charge of each bead $q$ to $-2e$, which is 11 times smaller than the combined charge of an 11-bp segment of DNA. In real dsDNA, each base pair of a dsDNA has a charge of $-2e$, therefore, the charge of each bead (11-bp segment of DNA) should be $-22e$. However, our goal is to just slightly perturb the system by adding the electrostatic interactions in such a way that the total contribution of the electrostatic interactions to the total potential energy of the system is small at $t = 0$ ns. In fact, in our case, the $U_{DH}$ contribution is about $(\frac{2}{22})^2 \sim 0.01$ of what it would be a full-strength Debye-Hückel potential was used.

Our simulations show that the explicit presence of long-range electrostatic interactions in the model polymer chain at $L_x/L_0 \approx 1.175$, $L_x/L_0 \approx 1.525$, and $L_x/L_0 \approx 1.85$ has little effect on the average values of the force (equation 4 of the main text) and the distributions (the probability density $\rho$) of relative bond lengths in the plateau regions (see "**un-s-electro**" in Table S1 and Figure S5).

##### Initial conformations

To check whether the simulation results depend on the initial conformation of the model polymer chain, we ran an additional set of simulations for the poly(dA-dT)-$\lambda$-DNA-poly(dG-dC) fragment at $L_x/L_0 \approx 1.175$, $L_x/L_0 \approx 1.525$, and $L_x/L_0 \approx 1.85$ from a conformation in which bond lengths alternate, i.e., slightly ($0.7r$) and highly stretched ($1.3r$), where $r$ is $1.175r_0$ (at $L_x/L_0 \approx 1.175$), $1.525r_0$ (at $L_x/L_0 \approx 1.525$), and $1.85r_0$ (at $L_x/L_0 \approx 1.85$). Our simulations show that the initial conformation of the model polymer chain has little effect on the average values of the force (equation 4 of the main text) and the distributions (the probability density $\rho$) of relative bond lengths $r/r_0$ in the plateau regions (see "**alter**" in Table S1 and Figure S5). Therefore, we conclude that the initial conformation of the model polymer chain has little effect on the force-extension curves and the distributions (the probability density $\rho$) of relative bond lengths $r/r_0$.

##### Integration time step

To check whether an integration time step of 100 fs is small enough, we ran an additional set of simulations with an integration time step of 50 fs for the poly(dA-dT)-$\lambda$-DNA-poly(dG-dC) fragment at $L_x/L_0 \approx 1.175$, $L_x/L_0 \approx 1.525$, and $L_x/L_0 \approx 1.85$ from conformations in which the beads are equally spaced. Our simulations show that reducing the integration time step has little effect on the average values of the force (equation 4 of the main text) and the distributions (the probability density $\rho$) of relative bond lengths $r/r_0$ in the plateau regions (see "**un-s**" and "**un-s-50fs**" in Table S1 and Figure S5). Therefore, we conclude that an integration time step of 100 fs is small enough.

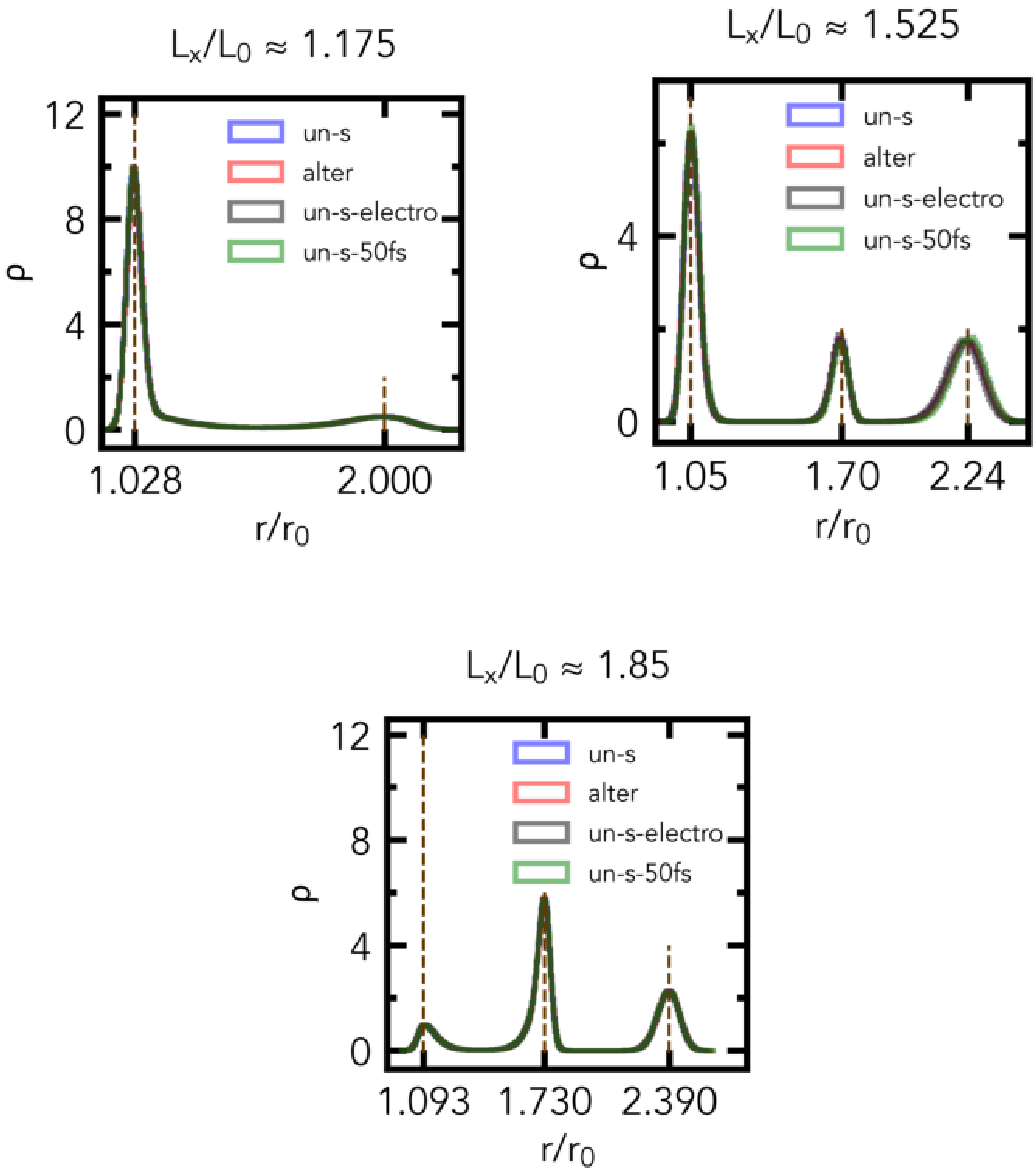


Figure S5: The distributions (the probability density $\rho$) of relative bond lengths $r/r_0$ in the plateau regions for the poly(dA-dT)-$\lambda$-DNA-poly(dG-dC) fragment at $L_x/L_0 \approx 1.175$, $L_x/L_0 \approx 1.525$, and $L_x/L_0 \approx 1.85$. The simulations start from (i) conformations of the chain in which the beads are equally spaced with an integration time step of 100 fs (**un-s**), (ii) conformations of the chain in which the beads are equally spaced with an integration time step of 50 fs (**un-s-50fs**), (iii) conformations of the chain in which the beads are equally spaced (the Debye-Hückel potential is added) with an integration time step of 100 fs (**un-s-electro**), (iv) conformations of the chain in which bond length alternate with an integration time step of 100 fs (**alter**). The relative bond lengths are recorded every 0.1 ns of the last 650 ns of the 700 ns simulation run.

**Table S1: Comparison of the average values of the force in the plateau regions for the poly(dA-dT)-$\lambda$-DNA-poly(dG-dC) fragment. The simulations start from (i) conformations of the chain in which the beads are equally spaced with an integration time step of 100 fs (un-s), (ii) conformations of the chain in which the beads are equally spaced with an integration time step of 50 fs (un-s-50fs), (iii) conformations of the chain in which the beads are equally spaced (the Debye-Hückel potential is added) with an integration time step of 100 fs (un-s-electro), (iv) conformations of the chain in which bond length alternate with an integration time step of 100 fs (alter). The values of the force (equation 4 of the main text) and relative extension $L_x/L_0$ are recorded at each integration time step, and then averaged over the last 650 ns (6.5 million values) of the 700 ns simulation run.**

| $L_x/L_0$ | un-s (pN) | alter (pN) | un-s-electro (pN) | un-s-50fs (pN) |
|---|---|---|---|---|
| 1.174 | 33.23 | 34.12 | 32.67 | 33.47 |
| 1.524 | 68.34 | 68.58 | 65.13 | 70.78 |
| 1.849 | 110.33 | 110.25 | 109.14 | 110.32 |
